# Nanopore sequencing for N1-methylpseudouridine in RNA reveals sequence-dependent discrimination of the modified nucleotide triphosphate during transcription

**DOI:** 10.1101/2022.06.03.494690

**Authors:** Aaron M. Fleming, Cynthia J. Burrows

## Abstract

Direct RNA sequencing with a commercial nanopore platform was used to sequence RNA containing uridine (U), pseudouridine (Ψ), or N1-methylpseudouridine (m^1^Ψ) generated by in vitro transcription (IVT). The base calling data as well as the ionic currents and dwell times for U, Ψ, or m^1^Ψ as they translocated through the helicase and nanopore proteins identified diagnostic signatures for Ψ and m^1^Ψ; however, the two modifications yielded similar patterns although both were different from U. Understanding the nanopore signatures for Ψ and m^1^Ψ enabled a running start T7 RNA polymerase assay to study how competing mixtures of UTP with ΨTP or m^1^ΨTP lead to nucleotide selection in all possible adjacent sequence contexts. For UTP vs. ΨTP, ΨTP was favorably incorporated in singly-modified contexts, while doubly-modified contexts found high yields of ΨTP insertion on the 5′ side and lower yields on the 3′ side. For UTP vs. m^1^ΨTP, UTP was favorably selected except in 5′-XA (X = U or m^1^Ψ) where the ratio was determined by their relative NTP concentrations. Experiments with chemically-modified triphosphates and DNA templates designed based on the structure of T7 RNA polymerase provide a model to explain the observations. These results may aid in future efforts that employ IVT to make therapeutic mRNAs with sub-stochiometric amounts of m^1^Ψ.

## Introduction

Inspection of native RNA across all phyla of life has identified >150 chemical modifications that include the addition of alkyl groups on the base and/or sugar, isomerization, sulfurization, oxidation, or reduction of the nucleobases (1-3). These chemical changes are found in tRNA, rRNA, mRNA, and small and large non-coding RNAs. The epitranscriptome refers to these modifications essential for the transcriptome’s functional relevancy for cellular processes. Additionally, chemical modification of RNA has found its way into clinical applications where the successes of therapeutic siRNAs and mRNA vaccines were largely achieved due to the site-specific chemical decorations of the polymer (4,5). Identification and quantification of RNA modifications have been pursued by nuclease and phosphatase digestion of target strands to nucleosides for LC-MS/MS analysis resulting in the complete loss of all sequence information (1). The development of nanopores as a third-generation sequencing platform has brought about many enabling advancements, one of which is the ability to directly sequence RNA. Out of the many benefits to direct RNA sequencing is the opportunity to sequence chemical modifications.

Nanopore sequencing on a commercial platform (Oxford Nanopore Technologies) is achieved by the utility of a helicase as a motor protein to deliver the RNA into a protein nanopore at a velocity that is ATP-dependent (Figure 1) (6). An electrophoretic force serves to guide the direction of the RNA 3′ to 5′ into the nanopore protein where a small central constriction zone exists. As the strand passes the constriction zone with a length of 5-nt of RNA (i.e., k-mer), the ionic current changes as a function of sequence (7). In recent iterations of base-calling software, the current vs. time traces are deconvoluted via a recurrent neural network (8). Exciting developments using nanopores have showcased direct RNA sequencing for modifications in tRNA, rRNA, mRNA, small/large RNAs, and viral RNAs from biological sources (7,9-15). The predominant native RNA modifications inspected include pseudouridine (Ψ), *N*^*6*^-methyladenonsine, and 2′-*O*-methyl nucleotides. Our work with Ψ demonstrated base calling error analysis combined with current and dwell time analysis minimizes the false discovery rate for modification detection in noisy nanopore data (12). Some challenges remain before this approach becomes a routine technique in the RNA researcher’s toolbox.

**Figure 1.**
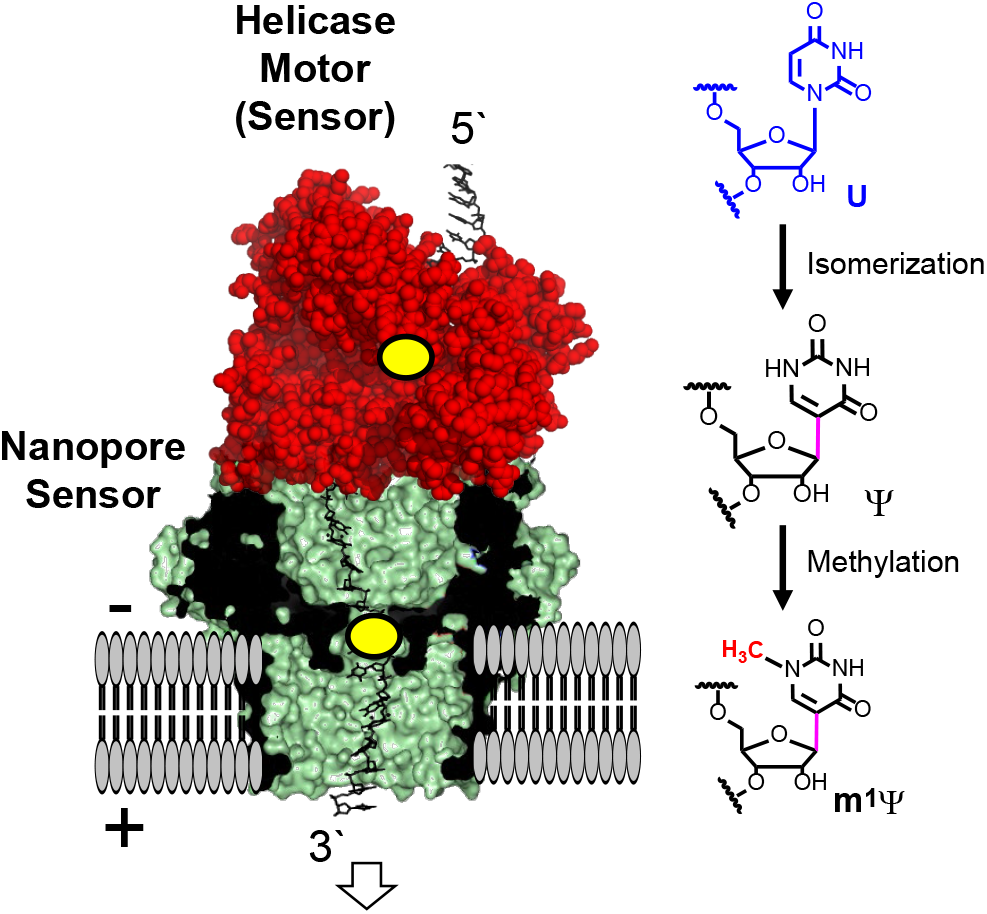
The Oxford Nanopore Technologies platform sequences U, Ψ, and m^1^Ψ directly in RNA.

In the present work, the commercial nanopore sequencer was employed to directly sequence N1-methylpseudouridine (m^1^Ψ) in RNA (Figure 1). This modification results from isomerization and methylation of the uridine nucleotide, and was first discovered in archaea tRNA by the McCloskey laboratory (16). Since then, this modification was found to be essential for the successful use of mRNA vaccines to defend against SARS-CoV-2 infections (5). These synthetic mRNA vaccines are produced by in vitro transcription (IVT), in which all U sites are converted to m^1^Ψ by feeding T7 RNA polymerase m^1^Ψ-nucleotide triphosphate (m^1^ΨTP) instead of UTP (5). Using the same approach for RNA production as the mRNA vaccines, synthetic RNAs with m^1^Ψ or Ψ were prepared and nanopore sequenced herein. The obtained data was inspected for changes in the ionic current, dwell time, and base calling profile between the modified RNAs and the U-containing RNA control strand of the same sequence. This analysis led to the design of new RNAs using the N1-ethyl or N1-propyl derivatives of Ψ to provide trending data regarding the nanopore signatures. These foundational results for the signals associated with Ψ and m^1^Ψ enabled a study of T7 RNA polymerase selection of the modified nucleotide triphosphate when the canonical UTP was present. All adjacent sequence contexts were studied. The nanopore sequencing found many unexpected sequence-context effects for canonical vs. non-canonical nucleotide triphosphate selection by the RNA polymerase. The present direct RNA sequencing via nanopores provided new insight into the clinically relevant modification m^1^Ψ, and how T7 RNA polymerase selects mixed competing NTPs during mRNA synthesis. These final results will be of critical importance for future manufacturing of therapeutic RNAs made by in vitro transcription that aim to have sub-stochiometric incorporation of modified nucleotides such as m^1^Ψ.

## Materials and Methods

### RNA synthesis by in vitro transcription

In vitro transcription was performed using the MEGAscript T7 transcription kit according to the manufacturer’s instructions. The duplex DNA templates for the IVT reactions were synthesized via commercial sources to have a T7 promoter for initiation of transcription and ended with a poly-A tail for sequencing library preparation (Figure S1). The IVT reactions were incubated overnight at 37 °C in a PCR thermocycler. After the overnight incubation, turbo DNase I treatment was performed on all samples at 37 °C, followed by purification using Quick Spin Columns for radiolabeled RNA purification. To install Ψ or m^1^Ψ, IVT was conducted in the presence of pseudouridine-5’-triphosphate (ΨTP) or m^1^ΨTP instead of UTP. Success in the synthesis of the RNA transcripts was verified by agarose gel electrophoresis by comparison to a ladder of known lengths. All experiments were conducted in duplicate.

### T7 RNA polymerase studies with mixed NTPs

Studies on NTP discrimination by T7 RNA polymerase were conducted using the MEGAscript T7 transcription kit with some changes to the manufacturer’s protocol as described. The NTP concentrations were 2 mM for ATP, GTP, and CTP, while the UTP and ΨTP or m^1^ΨTP (Trilink Biotechnologies) were 1 mM each to achieve a total concentration of U and its derivatives of 2 mM. To ensure the UTP and ΨTP or m^1^ΨTP were at a 1:1 ratio, the stock solution concentrations were determined using UV-vis spectroscopy with established extinction coefficients (UTP: λ_262 nm_ = 10,000 L mol^-1^cm^-1^; ΨTP: λ_262 nm_ = 7,550 L mol^-1^cm^-1^; m^1^ΨTP: λ_271 nm_ = 8,870 L mol^-1^cm^-1^). The IVT reactions were allowed to progress for 2 h at 37 °C before termination. Addition of turbo DNase I following the manufacturer’s protocol terminated the reaction. Further studies on T7 RNA polymerase NTP selection were conducted with N1-ethylpseudouridine triphosphate (e^1^ΨTP: λ_271 nm_ = 7,800 L mol^-1^cm^-1^) or N1-propylpseudouridine triphosphate (p^1^ΨTP: λ_271 nm_ = 8,900 L mol^-1^cm^-1^; Trilink Biotechnologies). All experiments were conducted in duplicate.

### Nanopore library preparation and sequencing

The poly-A tail containing RNAs generated by IVT were the input strands in the direct RNA sequencing kit (SQK-RNA002) from Oxford Nanopore Technologies (ONT). The protocol was followed without changes and the library prepared samples were directly used for sequencing. The samples were applied to the ONT Flongle™ flowcell running the R9.4.1 chemistry following the manufacturer’s protocol with an applied voltage of 180 mV. All sequencing experiments were conducted until all nanopores were no longer functional that generally yielded >15,000 reads.

### Data analysis

The ionic current vs. time traces in fast5 file format were base called using guppy v.6.0.7 to obtain the fastq sequencing read files used in the subsequent data analyses. The fastq files were aligned to the reference sequences using minimap2 ‘-ax map-ont -L’ to generate bam files (17). The bam file alignment statistics were determined with the flagstat function in Samtools and then the files were indexed with Samtools for visualization in Integrative Genomics Viewer (IGV) to obtain the base call information at the modification sites (18,19). The Nanopore-Psu, Nanocompore, and Tombo tools were used as described on their Github sites (7,11,20). The current and dwell time data were extracted, resquiggled to the reference, and analyzed using Nanopolish as described in the user manual (21). The data were plotted and analyzed in either python, Origin, or Excel for visualization.

## Results and Discussion

### Experimental setup

The RNA to address the performance of the nanopore sequencer on m^1^Ψ with comparisons to U and Ψ were synthesized by IVT, like the present approach for mRNA vaccine development (Figure S1) (5). The use of commercially available m^1^ΨTP or ΨTP allowed the generation of RNAs with 100% incorporation of the modified nucleotides. The RNA strands sequenced had the U, Ψ, or m^1^Ψ spaced >18 nts apart such that only one site of analysis interacted with the nanopore-helicase setup at a time. interacted A limitation to this approach is sequence contexts with the parent nucleotide U cannot be generated. The goal is to study T7 RNA polymerase activity in the presence of modified and canonical nucleotides to measure selectivity imposed by the enzyme; therefore, mixed sequence contexts with U and Ψ or m^1^Ψ is not an issue. All possible 5’ and 3’ adjacent sequence contexts flanking a single site and sites for double incorporation of the modification were designed into the duplex DNA template used for transcription of the RNA. The nanopore sequencer using the CsgG protein nanopore (v 9.4.1 flow cells) has a sensing window for RNA (i.e., k-mer) of ∼5 nt (7). The RNA sequences studied partially covered the sequence space for the nanopore k-mer; however, not all sequence combinations were studied.

### Base-calling error analysis

Nanopore sequencing was conducted on the RNA strands following standard protocols to generate current vs. time traces that were base called with Guppy (v 6.0.7), followed by alignment to the reference with minimap2, and visualization of the base calls was achieved with IGV. First, the base-calling error identified by the presence of calls for C, A, G, or insertion and deletions (indels) for the U, Ψ, and m^1^Ψ RNAs were quantified. In the data for sequences with two U, Ψ, or m^1^Ψ adjacent to one another, each of the two in the pair gave unique base call errors; thus, in the plots shown in Figure 2, the modified sites in the sequences are shown with the letters X and Y, where the data above the sequence are for the position labeled X. The U-containing RNA strands gave percent errors <10% for the sequence contexts studied (Figures 2A gray line). The percent error for each of the sequence contexts possessing Ψ was quantified and ordered from high to low (Figure 2A orange line) to find that the error ranged from nearly 100% down to ∼20% with a strong dependency on the sequence context. The percent base calling error for each m^1^Ψ-containing context was plotted (Figure 2A blue line). The data were not identical to those observed for Ψ, but they did show a similar trend. The base call analysis identifies modification of U to either m^1^Ψ or Ψ generates base called data with much higher error than U.

**Figure 2.**
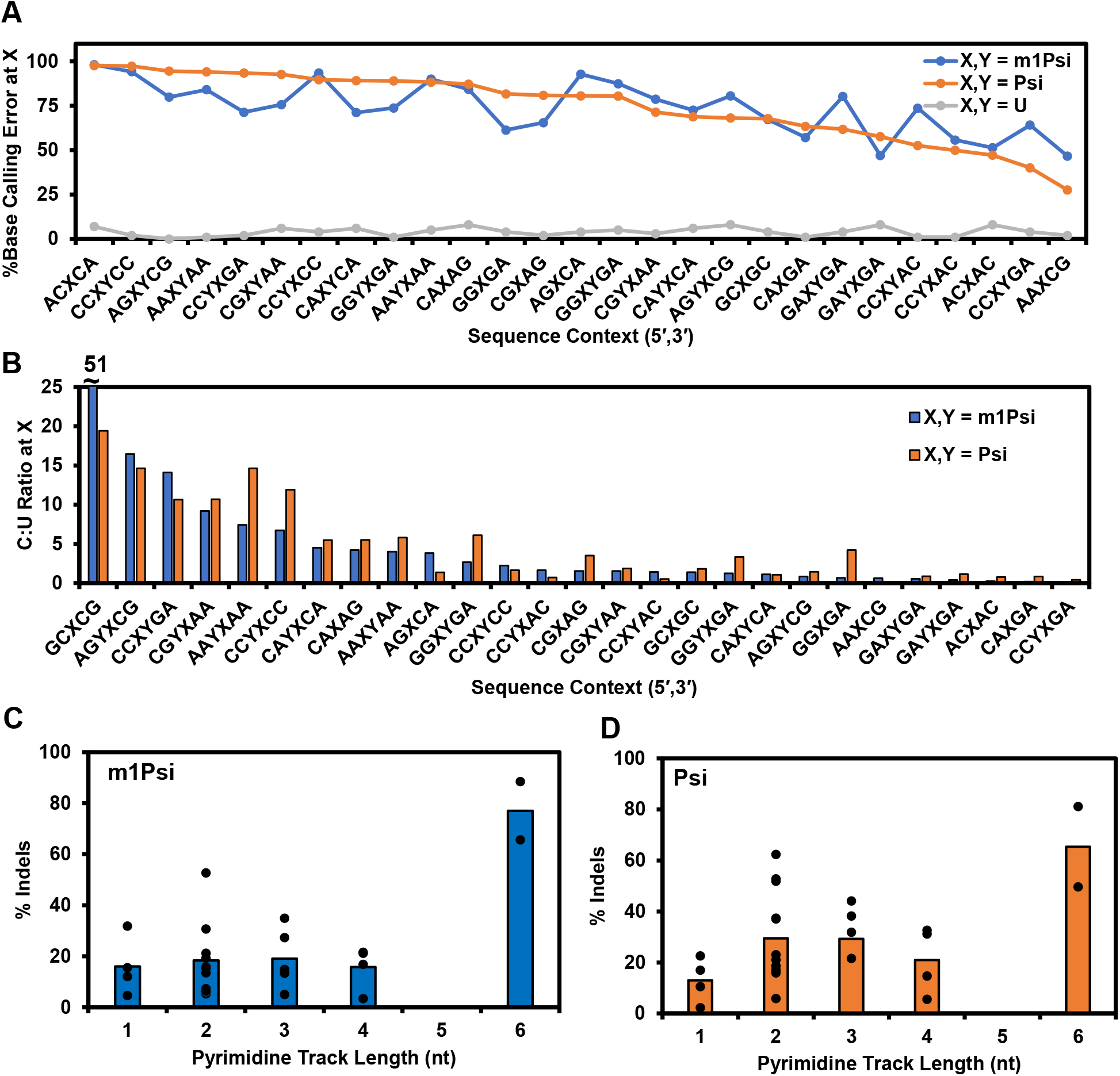
The presence of Ψ and m^1^Ψ yield sequence-dependent base-calling errors during direct RNA nanopore sequencing. (A) Plots of sequence context vs. percent total error, which include calls to A, C, G, and indels, for U (grey), Ψ (orange), and m^1^Ψ (blue). The values were generated from >1000 aligned reads. (B) A plot of sequence context vs. the C:U base calling ratio from the data in panel A for Ψ (orange) and m^1^Ψ (blue). The percent indels vs. pyrimidine track length from the panel A data for (C) m^1^Ψ and (D) Ψ. Some of the C:U base calling ratio data were previously reported by our laboratory (12) and have been reanalyzed for the present comparisons.

The base calling data were then inspected more closely to identify the neural network base caller error modes at the modified sites. Previous studies found Ψ is miscalled as a C (10-12,14). The C to U ratio for Ψ and m^1^Ψ in each context were compared to find they produced similar sequence dependency in the C relative to U calls (Figure 2B). Sequence-dependent trends in the base calling percent error or base call ratio were not observed. Next, the indel profiles for each context bearing Ψ or m^1^Ψ were quantified. The literature identified indels increase on the v 9.4.1 flow cell chemistry of the ONT platform in homopolymer sequence runs (22); with this in mind and knowing that the modifications can code like a C, the indel percentage was plotted against the length of the pyrimidine track flanking the site. These data do not enable the study of all possible sequence contexts, although when the pyrimidine track length was 6 nucleotides long, indel signatures for Ψ and m^1^Ψ increased. The increased frequency of indels observed in the reported data for 2-nt long tracks results from the over-representation of sequences with two modifications adjacent to one another in the sequence.

The key finding in the base calling analysis was that Ψ and m^1^Ψ cause changes to the raw data that the neural network base caller interprets to yield similar base-calling errors. Direct RNA nanopore sequencing for Ψ has been reported and tools developed to quantify the data to determine the extent of occupancy at modified sites (10-15). Many of these approaches rely on base calling features. The dominant and unique base calling signature for Ψ is to be called as a C (Figure 2B), which was found herein and previously (10,12,14,15). This approach in the analysis is subject to false positives when natural U→C variations exist in the RNA that can masquerade as a Ψ when inspecting base calling features exclusively; a solution to this challenge is described below (12).

The present work with m^1^Ψ identified its signature in the nanopore base calling data is like Ψ. This observation points to another challenge for direct RNA sequencing for modifications with nanopores, and that is different modifications can yield very similar signatures. After exploring the nanopore signatures for the >150 known modifications, the community might find, for example, that a particular U is modified in an unknown sample but the question of what the modification is will remain. Parallel biological and sequencing studies on native cells and those with suspected writer protein knockouts or knockdowns can aid in the identification of the modification (7,10). Alternatively, the use of low-throughput and targeted assays such as SCARLT or mass spectrometry sequencing will be needed to identify the modification, and even these will be challenged by modification isomers (e.g., m^5^C, m^4^C, and C_m_) (23,24).

### Ionic current level and dwell time analyses

Because the base calling data are derived from the current changes measured as the modification moves through the sensor, a question is whether Ψ and m^1^Ψ have similar current profiles. Another finding from our prior work was that Ψ impacted the helicase motor protein translocation kinetics to provide a second signal to follow the presence of the modification (12), which was confirmed in another study (25). We then inspected the current vs. time traces for U, Ψ, and m^1^Ψ to measure how they differ regarding the residual current levels and dwell times when the subject sites were in the nanopore, and the residual current levels and dwell times when the sites were in the helicase active site.

In Figure 3B, a stacked plot of current levels for either Ψ or m^1^Ψ relative to U obtained when they are in the nanopore sensor was generated using the Tombo tool (20). All other Tombo-generated plots are shown in Figure S2. This current-level analysis illustrates a very important point regarding nanopore sequencing for modifications. The nanopore constriction houses ∼5 nts of RNA called a k-mer and the change in current as a result of the modification does not have to occur when it is positioned in the center of the k-mer; as these examples show, the largest deviation in current occurred when the C 5′ to the modification was centered in the k-mer. As a consequence, each sequence context containing U or either of the two modifications was first analyzed visually with Tombo and statistically with Nanocompore (7) to locate the sites of the largest current deviation in the nanopore and dwell time deviation in the helicase for the plots provided. To avoid confusion, the k-mer sequences in all plots have the modification site in the center, while the actual k-mers analyzed are provided in the figure legend. This analysis was restricted to sequence contexts that had one modification because those with two generally induced current and dwell time changes that occurred at multiple locations (Figure S2). Single Ψ and m^1^Ψ modifications were analyzed in all possible adjacent sequence contexts that do not include the parent nucleotide U (9 contexts), and the 5′-GXC context was studied in two different 5-nt k-mers (5′-AGXCA and 5′-CGXCG where X = U, Ψ, or m^1^Ψ).

**Figure 3.**
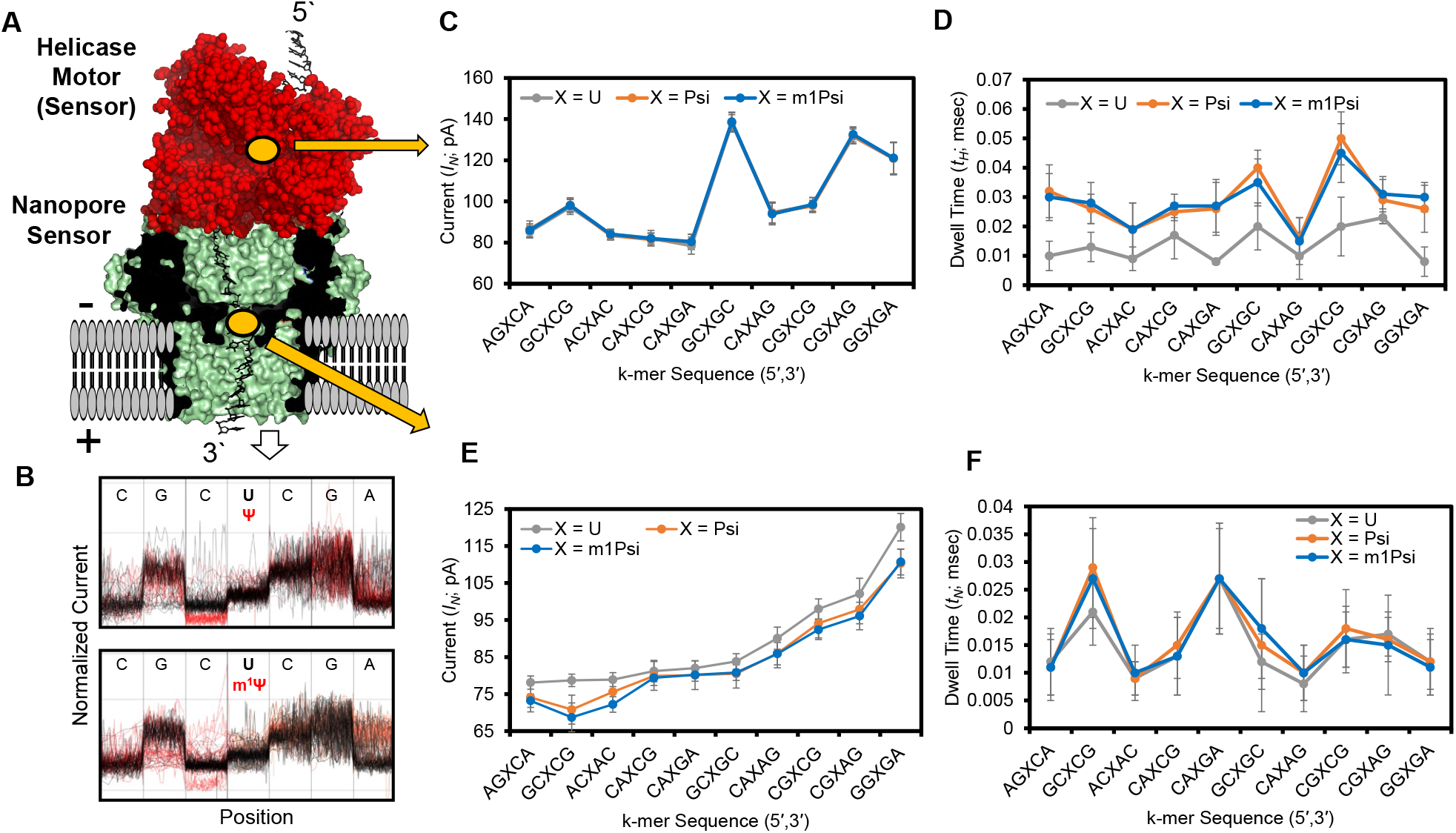
Analysis of currents and dwell times as sites of single U, Ψ, or m^1^Ψ resided in the nanopore or helicase sensors. (A) A structural view of the nanopore sequencer set up. (B) Comparative current level analysis for U vs. Ψ (top) or m^1^Ψ (bottom) generated by Tombo. These plots illustrate that the maximal current level change does not always occur when the modification is centered in the central constriction zone of the CsgG protein. Additionally, Tombo plots compress the time domain on the x-axis for visualization; therefore, the x-axis or position does not reflect the true residence times. (C) Comparative plots of the helicase current levels (*I*_*H*_) and (D) dwell times (*t*_*H*_) for the sequence context studied. (E) Plots for comparison of the nanopore current levels (*I*_*N*_) and dwell times (*t*_*N*_) for the sequence context studied. The position of the maximal differences between U and the modifications was derived from Tombo and Nanocompore analysis. The actual k-mers analyzed for the nanopore sensor *I*_*N*_ and *t*_*N*_ values are provided in parentheses written 5′ to 3′ where X = U, Ψ, or m^1^Ψ: AGXCA (GXCAC), GCXCG (GCXCG), ACXAC (XACCA), CAXCG (ACAXC), CAXGA (GCAXG), GCXGC (GCXGC), CAXAG (CAXAG), CGXCG (CGXCG), CGXAG (ACGXA), and GGXGA (XGACA).

The current and dwell values were obtained from Nanopolish resquiggled data (i.e., aligning base calls with current levels). Previously it was found that resquiggling with all available software (Tombo or Nanopolish) can filter out data and lead to biases in ways that are not well understood (10), which was confirmed herein (Figure S3); however, no other tools are available for this type of analysis. With this known issue in mind, we proceeded with caution and avoid overarching claims regarding the data. The U, Ψ, or m^1^Ψ first interact with the sequencer in the helicase where the residual current (*I*_*H*_) and/or average dwell time (*t*_*H*_) can be impacted. The *I*_*H*_ values for U, Ψ, and m^1^Ψ for each sequence context were nearly identical (Figure 3C). This is consistent with the fact that the source of the current deviations is in the nanopore, not the helicase. When the *t*_*H*_ values were inspected for U, Ψ, or m^1^Ψ in the helicase, the times were longer for Ψ and m^1^Ψ relative to U (Figure 3D). This is consistent with our previous finding regarding the helicase stalling on Ψ (12). A comparison of the average *t*_*H*_ for Ψ and m^1^Ψ for the contexts studied found they are similar (Figure 3D).

Next, the interrogated site interacts with the nanopore sensor in which the residual currents (*I*_*N*_) and average dwell times (*t*_*N*_) were measured. The *I*_*N*_ values for the k-mers found changes between U vs. Ψ or m^1^Ψ (Figure 3E). All U-containing k-mers had greater *I*_*N*_ values relative to Ψ or m^1^Ψ in the same context, which shows the modifications are more blocking to the current. The *I*_*N*_ values between Ψ and m^1^Ψ were similar. The average *t*_*N*_ values for the three nucleotides in each sequence context gave small differences (Figure 3F), but they were less than those observed for the modifications in the helicase. The key regions Ψ and m^1^Ψ impact the recorded data are in the dwell time while situated in the helicase active site and in current level while located in the nanopore central constriction zone. The changes in *I*_*H*_, *t*_*H*_, *I*_*N*_, and *t*_*N*_ for each of the singly-modified k-mers were inspected for correlations with the base calling error, C:U ratio, and indel frequency using a Spearman’s rank-order correlation test. The analysis found for Ψ the base calling error and C:U ratio correlated with *I*_*N*_ (ρ = 0.88 and ρ = 0.84) but not with the other physical values measured; in contrast, indel frequency for Ψ did not correlate with any parameter measured (Figure S4). The correlation analysis for m^1^Ψ, found *I*_*N*_ correlated with base calling error (ρ = 0.61), while *I*_*N*_ and *t*_*N*_ correlated with the C:U ratio (ρ = 0.58 and ρ = -0.61; Figure S4). These correlations are not surprising; however, they led to a testable hypothesis as described below.

The nanopore sequencer is unique because proteins are used in a setup to generate ionic current vs. time traces for DNA or RNA as they pass through the platform under an electrophoretic force. The nanopore protein and helicase motor can function differently on RNA with chemical modifications compared to the all-canonical strands to provide signatures for the presence of these changes. The U modifications Ψ and m^1^Ψ impacted the dwell time in the helicase sensor and the current level in the nanopore sensor to provide more data for confident calls of the modifications. Inspection of the current vs. time data can be beneficial when conducting de novo sequencing of RNA for modifications. Using Ψ and m^1^Ψ as an example, they are base called like C nucleotides; therefore, natural U→C variations would exist as false positives when looking for these modifications in base calling analysis. An approach to differentiate modifications from sequence variations is to look at the helicase dwell time *t*_*H*_, as we previously proposed (12).

### Study of N1-methyl-, N1-ethyl-, and N1-propyl-Ψ derivatives

Two main motivations led us to study the N1-methyl, N1-ethyl, and N1-propyl derivatives of Ψ (i.e., m^1^Ψ, e^1^Ψ, and p^1^Ψ; Figure 4A) with the nanopore sequencer. First, the previous observation that the nanopore current values *I*_*N*_ showed a correlation with base calling features for Ψ and m^1^Ψ. This led us to wonder whether systematically increasing the alkyl size would impact the *I*_*N*_ value leading to a trending change in the base calling features. Said another way, can systematic chemical changes to the base result in expected changes in how the neural network base caller identifies bases. Second, a patent filed for use of nucleotide modifications for mRNA vaccines has included claims with methyl, ethyl, and propyl derivatives (26), and therefore, nanopore sequencing of these modifications may have relevance in the future. Moreover, nucleotide triphosphates for N1-ethyl and N1-propylpseudouridine are commercially available.

**Figure 4.**
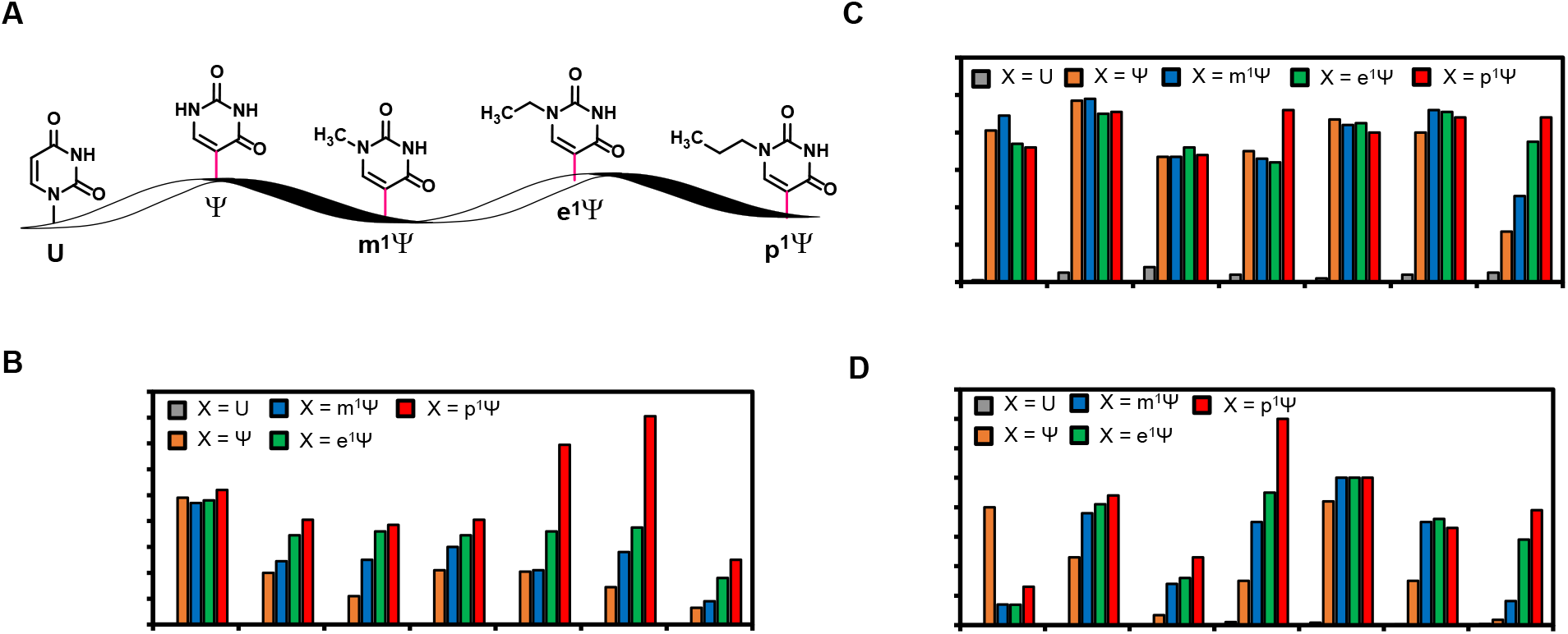
Nanopore sequence analysis of the N1-alkyl derivatives of Ψ. (A) The structures of U, Ψ, m^1^Ψ, e^1^Ψ, and p^1^Ψ. (B) The difference between the U *I*_*N*_ value and the N1-alkyl Ψ *I*_*N*_ values (Δ*I*_*N*_ = U *I*_*N*_ – X *I*_*N*_; where X = Ψ, m^1^Ψ, e^1^Ψ, or p^1^Ψ). (C) The percent base calling error, and (D) the C:U base call ratio for each of the nucleotides studied. These studies were conducted on seven different sequence contexts where > 500 single-molecule measurements from two replicates provided the average values reported.

A subset of singly-modified sequence contexts in RNA was synthesized by IVT, libraries were prepared and then sequenced by the nanopore. The data were resquiggled with Nanopolish to then find the values *I*_*N*_, *t*_*N*_, *I*_*H*_, and *t*_*H*_ for each context and N1-alkyl Ψ derivative. Noteworthy, as the alkyl group size increased the percentage of reads lost during resquiggling with Nanopolish increased (Figure S3). Figure 4B provides a plot of the difference in *I*_*N*_ values between the U-containing RNA with each N1-modified Ψ studied (i.e., Δ*I*_*N*_ = [U *I*_*N*_ - modified Ψ *I*_*N*_]). All current and dwell time average values with the associated errors are provided and described in Figure S5. Out of the seven sequence contexts studied, six showed a trend in the Δ*I*_*N*_ values with the size of the alkyl group. The exception was a modification in the context 5′-GGXGA that produced a large deviation from U to Ψ, for which the alkyl group size did not change this already large difference. Analysis of the percent base calling error for the modifications in the different sequence contexts did not identify a visible trend, except the k-mer 5′-CAXAG. This standout context is the one that showed the least base calling error for Ψ, and therefore, there was room for an increased change to be observed for the increasing alkyl group size on N1. Lastly, the C:U base calling ratio was inspected, and in all but one case, the value increased going from Ψ to the N1-alkyl derivatives, and the change varied with sequence context. The context 5′-GGXGA was the one that gave a decrease in the C:U ratio for Ψ and its derivatives, and the reason for this is not known.

These data identify the series Ψ, m^1^Ψ, e^1^Ψ, and p^1^Ψ generate nanopore-derived base calling error differences from U that can be followed to identify their presence in an RNA strand. Additionally, the currents and dwell times for the N1-alkyl Ψ derivatives do change (Figure S5), but the data are subject to loss during the resquiggling process resulting in a low confidence in the values obtained (Figure S3). Hopefully, future efforts on the computational side of nanopore sequencing analysis will find a solution to this limitation. These data may assist future efforts that use selective chemistry to add chemical reporter groups to target modifications for nanopore sequencing (27). This pilot study provides promise for this approach but reminds us that sequence context will influence the results in unexpected ways.

### T7 RNA polymerase NTP selection for mixtures of UTP with ΨTP or m^1^ΨTP

In the final set of studies, we used our knowledge of nanopore sequencing signatures for Ψ or m^1^Ψ to explore NTP selection by T7 RNA polymerase during IVT. Currently, mRNA vaccines are produced by T7 RNA polymerase mediated transcription, in which all the U nucleotides are completely replaced with m^1^Ψ (5). A prior study found the partial replacement of the canonical nucleotides with modified forms (m^5^C and s^2^U) in therapeutic mRNAs generated by IVT can be effective (28); however, the studies did not know whether there existed sequences that T7 RNA polymerase favored or disfavored insertion of the non-canonical nucleotide triphosphates. Therefore, we addressed this question for UTP vs. ΨTP or m^1^ΨTP mixtures during T7 RNA polymerase synthesis of an mRNA. The duplex DNA template provided coding potential to interrogate all possible immediate sequences contexts in singly-modified and doubly-modified contexts.

The RNA polymerase evaluation for NTP selection was conducted with a one-to-one ratio of UTP to ΨTP or UTP to m^1^ΨTP. To ensure the competing NTPs were mixed in a one-to-one ratio, the stock solution concentrations were determined by UV-vis spectroscopy using extinction coefficients provided by the manufacturer of the nucleotides. The RNA strands were synthesized and then sequenced with the nanopore. The synthesized RNAs were analyzed by agarose gel analysis to confirm the full-length extension of the template (Figure S6). Quantification of the Ψ or m^1^Ψ insertion was achieved using a recently published tool, Nanopore-Psu (11), which was trained on base calling data to use the associated errors for prediction of the modification status at each site. To ensure the best accuracy in quantification, and because the tool was designed for Ψ, not m^1^Ψ, we used the aligned nanopore reads to mix them in known ratios of U with Ψ or m^1^Ψ to develop calibration curves (Figure S7). This approach allowed the repurposing of Nanopore-Psu for occupancy prediction of another U modification (i.e., m^1^Ψ) in the sequence contexts studied.

The U vs. Ψ competition for the T7 polymerase active site in singly-modified contexts found Ψ was favorably installed with a 60-90% yield (Figure 5A). In the same competition but for sites in which the template DNA strand codes for insertion of two adjacent U/Ψ residues, something very interesting was found. In each context, the 5′ site favored ΨTP insertion (>50%) while the 3′ site gave a >2-fold reduction in ΨTP insertion (Figure 5B). This suggests insertion of the first modified nucleotide influences the insertion of the second.

**Figure 5.**
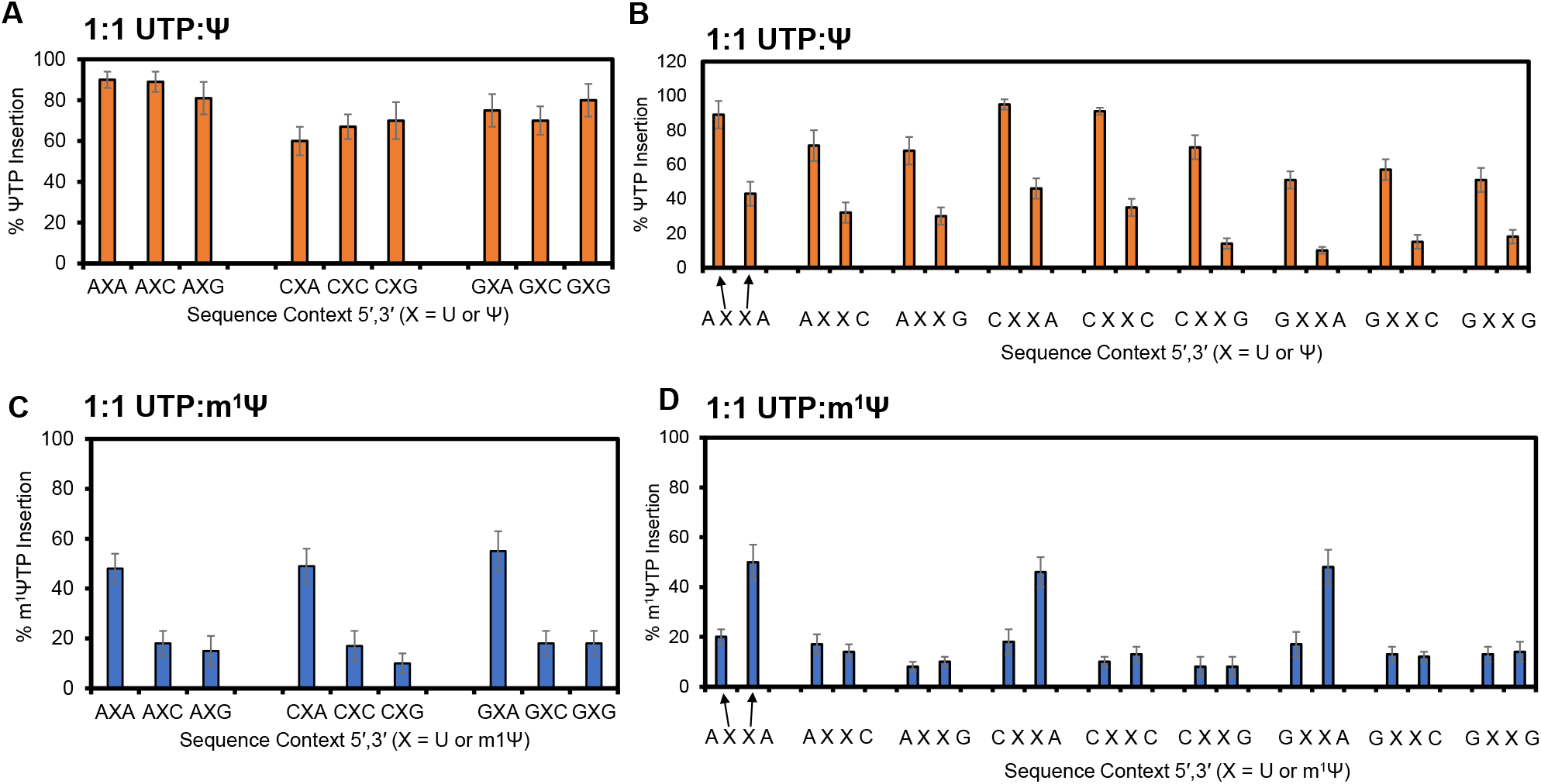
Yields of ΨTP or m^1^ΨTP incorporation when competed with UTP for insertion and elongation by T7 RNA polymerase. Percent insertion yields for ΨTP in (A) singly-modified or (B) doubly modified sequence contexts. Percent insertion yields for m^1^ΨTP in (C) singly-modified or (D) doubly-modified sequence contexts. The yields were determined via direct RNA sequencing with the commercial nanopore platform and the base calling data were analyzed with the published tool Nanopore-Psu (11).

Nanopore sequencing results from the competition between UTP and m^1^ΨTP for T7 RNA polymerase incorporation were even more different. In sites where the template codes for a single modification, UTP was installed with >80% yield with one exception; when the template DNA coded for ATP insertion 3′ to the modified nucleotide, T7 RNA polymerase inserted UTP and m^1^ΨTP with the frequency at which they were mixed in solution (i.e., 1:1; Figure 5C). Last, sites that could incorporate two UTP or m^1^ΨTP adjacent to one another found UTP was the major triphosphate incorporated except when there was a 3′ ATP inserted in the strand, which gave a nearly 1:1 ratio of the two competing pyrimidines being inserted (Figure 5D). The ability to directly sequence RNA with nanopores and quantitatively call modifications led to the discovery of the sequence-dependent bias that T7 RNA polymerase has for the selection of UTP vs. ΨTP or m^1^ΨTP.

Using the Nanopore-Psu tool to analyze direct RNA nanopore sequencing data, we studied T7 RNA polymerase selection of competing NTPs in many sequence contexts in a running start assay (Figure 6A). To address these results a better understanding of T7 RNA polymerase is needed. The phage RNA polymerase T7 is a single subunit DNA-dependent RNA polymerase, it does not have an exonuclease domain for proofreading, and with these limitations, the polymerase maintains high fidelity transcription (1 error in 10^4^ NTPs polymerized) (29). How T7 RNA polymerase maintains high fidelity RNA synthesis has been addressed by x-ray crystallography and computational studies (30-34). Experimental work identified that T7 RNA polymerase easily accepts modified NTPs (35), NTP elongation kinetics are influenced by steric factors (36), and NTP selection is impacted in solutions where the dielectric constant and water activity have been altered with PEG-200 (37). The key points in discrimination of the incoming NTP are to ensure the exclusion of dNTPs by checking the presence of the 2′-OH group, and that the NTP and templating DNA nucleotide form a viable base pair to maintain the fidelity of the mRNA code. Why does T7 RNA polymerase discriminate between UTP vs. ΨTP or m^1^ΨTP, of which can both form viable base pairs with the dA nucleotide in the template DNA strand?

**Figure 6.**
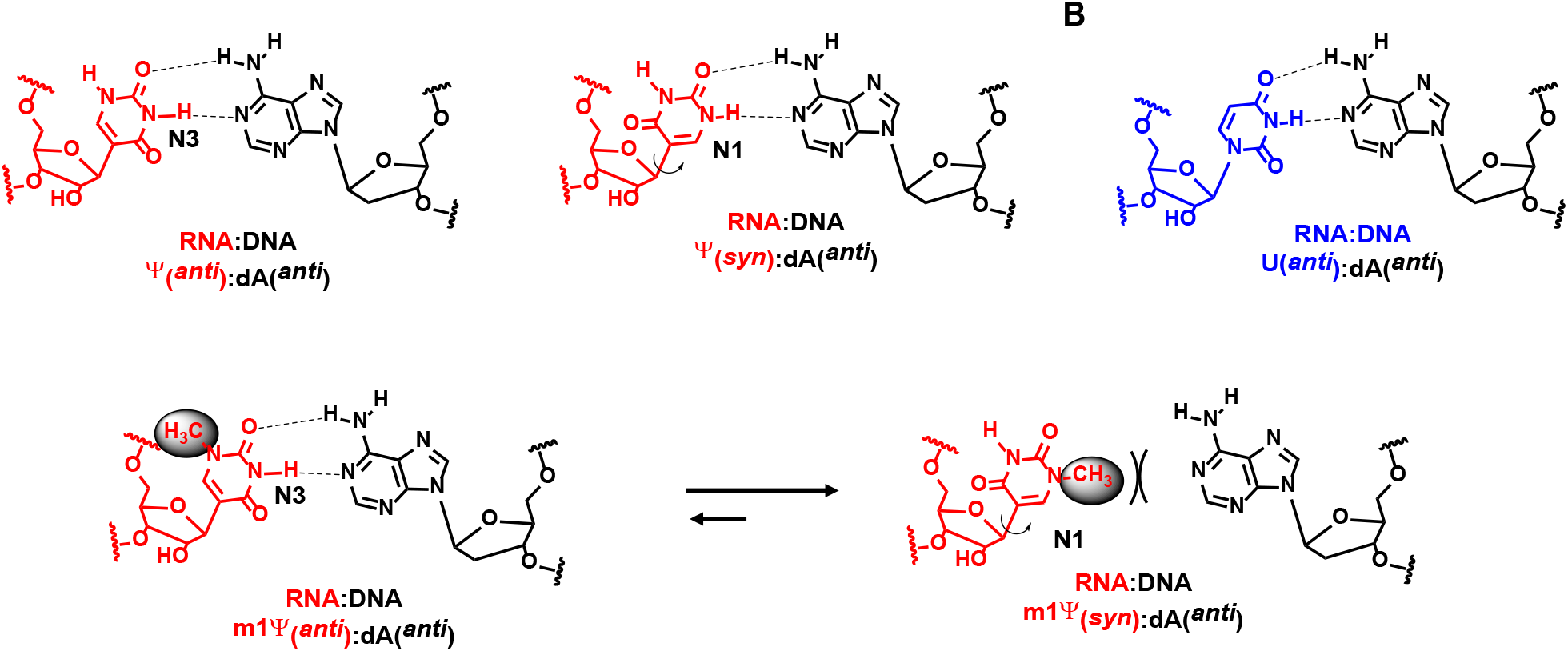
Base pairs formed by (A) Ψ, (B) U, and (C) m^1^Ψ in RNA with a dA nucleotide in a template DNA strand.

### T7 RNA polymerase selectivity for ΨTP when competing with UTP

When UTP competed with ΨTP for incorporation in singly modified sites, the yield of ΨTP incorporation was >60% (Figure 5A). We hypothesize the reason for the favorability of ΨTP incorporation is the greater freedom for *syn* and *anti*-glycosidic bond angles of the base relative to the sugar in this C-nucleoside that was found to favor the *syn* conformation (38). Both conformations of ΨTP display a face to the dA nucleotide that can form two hydrogen bonds with similar base-pair shapes (Figure 6A). Thus, ΨTP is favorability inserted because both base conformations *syn* or *anti* relative to the sugar yield viable dA base pairs, whereas UTP can only base pair with dA in the *anti* conformation (Figure 6B). This claim is supported by modeling work on T7 RNA polymerase that found phosphodiester bond formation for polymerization can only occur when the base pair has the right size and shape (34). When the template DNA strand had two adjacent dA nucleotides to direct insertion of the competing NTPs, ΨTP was favored for incorporation on the 5′ site but was disfavored by >2-fold for the 3′ site in all sequence contexts. A possible explanation for this observation comes back to the *syn*-and *anti*-glycosidic bond conformations for Ψ. Installation of a *syn* Ψ at the 5′ site based on these data disfavors a second non-canonical nucleotide from being incorporated at the 3′ site. The 5′ syn Ψ lacks a carbonyl group in the minor groove that is known to be important in DNA polymerases for progression (39). Detailed structural studies will need to be conducted to address this structural hypothesis for two adjacent Ψ incorporations by T7 RNA polymerase.

### T7 RNA polymerase selectivity for UTP competed with m^1^ΨTP

Considering m^1^ΨTP vs. UTP for incorporation by T7 RNA polymerase, the findings differed considerably compared to competitions with ΨTP. In the sequence contexts that coded for a single U/m^1^Ψ, UTP was favorably selected by >4-fold (Figure 5B). The sequence context exceptions were those with a 3′ A in the RNA that gave U and m^1^Ψ incorporation at a 1:1 ratio reflected by their concentrations in solution. There are two mysteries regarding these observations. (1) Why does UTP outcompete m^1^ΨTP in all sequence contexts except one? (2) Why does a 3’ A in the RNA result in higher yields for the modified nucleotide triphosphate?

The Chow laboratory used NOE NMR measurements to report on the *syn* vs. *anti* conformational equilibrium for m^1^Ψ in the nucleoside context (38). They found m^1^Ψ had a greater preference for the *syn* conformation than Ψ. Unlike Ψ, m^1^Ψ *syn* cannot base pair with dA in the DNA template strand (Figure 6C); therefore, the dominant glycosidic bond conformation for this modification is not compatible with T7 RNA polymerase to catalyze phosphodiester bond formation. This results in UTP outcompeting m^1^Ψ for incorporation and elongation by this DNA-dependent RNA polymerase.

Regarding the 3′ A effect where m^1^ΨTP and UTP were selected based on their solution concentrations, the structures for the steps of NTP selection and polymerization by T7 RNA polymerase offer a clue that can be tested (30-34). The active site of T7 RNA polymerase is maintained in an open conformation by the O-, O’-helix during NTP selection that closes to generate the contacts needed for NTP discrimination and phosphodiester bond formation (30,33). Computational studies suggest closing of the O-, O’-helix is thermally regulated and that the open conformation provides space for NTPs to sample glycosidic bond conformations to yield viable base pairs for polymerization in the growing mRNA strand (32). In the open and closed conformations, the DNA base pair 3′ to the templating nucleotide is still formed and a pi stack between F644 of the O, O’-helix, and the template DNA nucleotide exists (Figure 7A). In this case, the pi stack would be with a dT on the template DNA strand to form the weakest interactions possible, a pi stack with a pyrimidine in a two-hydrogen bond base pair. This may allow greater sampling of the open and closed states of the O-, O’-helix providing space and time for m^1^ΨTP to find the *anti* conformation to base pair with the template dA (Figure 6C).

**Figure 7.**
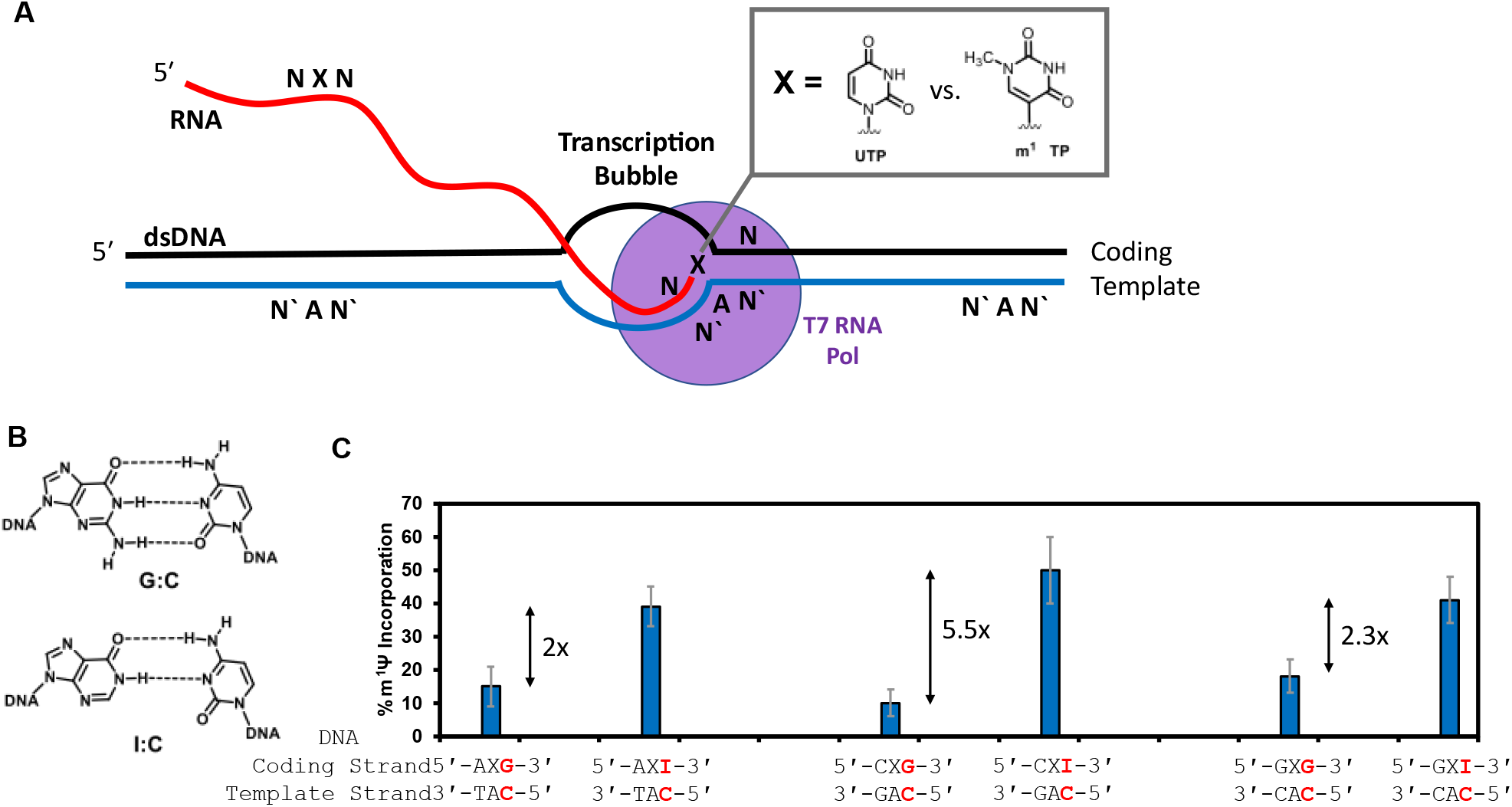
The T7 RNA polymerase structure during transcription provides a testable hypothesis why UTP and m^1^ΨTP are selected equally when an A is 3’ to the site in RNA. (A) A cartoon model of transcription by T7 RNA polymerase focusing on the base pairs. (B). Structures for the base pairs dG:dC and dI:dC. (C) Yields for m^1^ΨTP incorporation when a dI:dC base-pair follows the site of competition to give yields similar to the situation with a dA:dT base pair at that position.

To study this polymerase-substrate active site structural proposal, a duplex DNA was designed, synthesized, and used for IVT that had a DNA templating dT nucleotide for the UTP vs. m^1^ΨTP competition followed by a dC. The change was in the coding DNA strand where the native dG that forms three hydrogen bonds with dC was replaced with 2’-deoxyinosine (dI) for pairing with two-hydrogen bonds with the template dC. This situation would have a weaker base pair (2 hydrogen bonds) and pi stacking (dC) interaction with F644 but direct the insertion of a G 3′ to the U/m^1^Ψ competition site (Figure 7B). The nanopore sequence analysis found in the weakened 3′ base pair system in comparison to the dG:dC base pair the incorporation of m^1^Ψ increased by nearly >3-fold (Figure 7C). These results support the weaker protein-DNA interaction and DNA base pair on the 3′ side of a site for insertion of U or m^1^Ψ impact the NTP insertion and elongation yields. The stronger interactions (base pairing and pi stacking) favor canonical NTP insertion, while the weaker interactions allow non-canonical NTPs that must sample glycosidic bond angles for proper base pairing to compete. Controls were conducted with the dI:dC base pair on the 5′ side of the templating dA in the duplex DNA to find no effect on the U/m^1^Ψ insertion and elongation ratio (Figure S8). When the dA:dT is swapped to have the dA in the template DNA strand, the pi stack is stronger and UTP outcompetes m^1^ΨTP as was studied in the doubly-modified sequence contexts (Figure 5D).

As a final study regarding the T7 RNA polymerase competition assay, UTP was mixed in equal ratios with either e^1^ΨTP or p^1^ΨTP during IVT. The RNAs made were then directly sequenced and the percentage of modification insertion was quantified via the base calling analysis using Nanopore-Psu (11). Only singly-modified sequence contexts were analyzed. The analysis found as the alkyl group length increased at N1 of pseudouridine, the incorporation yield ratio for the modified NTP increased (Figure S9). The yields ratios were close to 1:1 for UTP vs. p^1^ΨTP. Prior work with long alkyl groups attached to the uridine nucleoside found them to adopt equal populations of *syn* and *anti* conformations (38). These results are of interest in situations where RNA is to be synthesized by IVT that would like a balanced distribution of U nucleotides with N1-alkylpseudouridine nucleotides.

## Conclusion

Direct RNA sequencing with nanopores can yield characteristic signatures for modifications that have been demonstrated for many modifications such as Ψ (10-13,15). In the present work, RNA containing either U, Ψ, or m^1^Ψ in all adjacent sequence contexts was generated by IVT and then sequenced with the commercial nanopore platform. The base calling data obtained identified unique signatures for the two U modifications that can be used to predict their presence in RNA. Using available computational tools, the occupancy of Ψ or m^1^Ψ can be predicted, as well. The present studies used the tool Nanopore-Psu first developed to predict Ψ occupancy in RNA (11) in the base called data for the analysis of m^1^Ψ. Further, we found the Ψ and m^1^Ψ relative to U yielded similar changes to the ionic currents when located in the nanopore sensor and slowed the dwell times when the modifications pass through the active site of the helicase motor protein. In the big picture, this analysis demonstrated different RNA modifications on the same nucleotide can yield similar signatures. This illustration should bring caution to RNA researchers using nanopore sequencing to conduct de novo analysis for chemical modifications. A site of modification could be found but the identity could remain questionable.

Understanding the nanopore sequencing data for Ψ or m^1^Ψ enabled a running start T7 RNA polymerase assay to be conducted to compete them (ΨTP or m^1^ΨTP) against UTP during transcription. The nanopore sequencing data for the RNAs found ΨTP outcompeted UTP in all singly-modified contexts that we hypothesize results from *syn* and *anti* Ψ pairing equally well with dA in the template DNA strand. In doubly-modified contexts, ΨTP was favored on the 5′ side and disfavored on the 3′ side that we propose results from the unfavourability of incorporating two *syn* pyrimidines in a growing mRNA chain. In contrast, UTP outcompeted m^1^ΨTP in all contexts except when a 3′ A occurred after the modification site in the RNA, where the yield ratios were the same as the NTP concentration ratio in the solution. A model for the results is proposed based on the prior finding that m^1^Ψ strongly favors the *syn* conformation and cannot pair with dA in the *anti* conformation allowing UTP to outcompete the modification (38). The exception is when there is an A coded for in the RNA sequence 3′ to the modification, which based on the solved structures for the polymerase (30,33) provides a more flexible active site to allow m^1^ΨTP to adopt the *anti* conformation to pair with the templating dA. The present results identify that when mRNAs are synthesized with a mixture of U and m^1^Ψ, their NTP concentrations will not be a good predictor of the extent of modification in the RNA because T7 RNA polymerase will function as a gatekeeper for the insertion yield of the NTPs in different sequence contexts. This information will be needed in future situations where therapeutic mRNAs are produced with sub-stoichiometric levels of m^1^Ψ, which has been successfully conducted with other RNA modifications (28).

## Data Availability

The data are available upon request.

## Supplementary Data

Supplementary data are available at XYZ.

## Acknowledgments

The National Institutes of Health provided financial support for this project (R01 GM093099). Oligonucleotide synthesis was provided by the University of Utah Health Sciences Core facilities that are supported in part by a National Cancer Institute Cancer Center Support grant (P30 CA042014).

