## Supplementary Data for "Nanopore sequencing for N1-methylpseudouridine in RNA reveals sequence-dependent discrimination of the modified nucleotide triphosphate during transcription"

| Item | Page |
|---|---|
| **Figure S1.** Sequences for the DNA templates used in the synthesis of the RNAs. | **S2** |
| **Figure S2.** Tombo analysis to compare the current levels between U and Ψ or U and $m^1Ψ$. | **S3** |
| **Figure S3.** Percent reads recovered after Nanopolish resquiggling. | **S9** |
| **Figure S4.** Spearman's rank-order correlation test on base call error vs. current levels and dwell times. | **S10** |
| **Figure S5.** Additional nanopore current and dwell data for the N1-alkylpseudouridine derivatives. | **S11** |
| **Figure S6.** Full-length extension evaluated by agarose gel electrophoresis. | **S12** |
| **Figure S7.** Calibration curves to quantify Ψ or $m^1Ψ$ developed with Nanopore-Psu. | **S13** |
| **Figure S8.** Controls with a dI:dC base pair on the 5′ side of the T7 RNA polymerase competition site. | **S14** |
| **Figure S9.** T7 RNA polymerase studies that competed UTP vs. $e^1Ψ$TP or $p^1Ψ$TP for insertion and elongation. | **S15** |

**Figure S1.** Sequences for the DNA templates used in the synthesis of the RNAs.

Coding Strand Sequence 1

5′-
AAGCTAATACGACTCACTATAGGAGCACAGGACCAGAC**GCTCG**ACAGAGCCGAAGCACAG
CAGACCAGAC**CTTCC**AGAAGACGAGACCA**ACTA**CCAGAAGCCGAAGCACAGACGA**AATTA**
**G**CCAGACGGACAACAGCAGAGACCGAAG**CATGAG**CAGACACGCAGCGACAGAGCAGCA**G**
**GTTGA**GGACCAGCAGGACAACAGGAGC**CTTAC**ACAGACGCGAACAACACAGA**CCTTG**AGC
CAGAGACAGACAC**AGATTGA**CACGAAGCGCAAGCAACCGCAGAA**CGTTAG**CACGAGCGG
CAACAGAGACGA**AGTTCG**GCAAGCACGCAGGCAACAGGACACGACGAGCGACC**ATTCA**G
AGACGACAGACACGCAGGCAGAGCACAAGCAAAACAAAAAAAAAAAA

Coding Strand Sequence 2

5′-
AAGCTAATACGACTCACTATAGGAGCAGCACGAGACGA**GGTGA**CACGACAGAGAGCGGAC
GC**AGTCA**CGACCGACGAACACGCA**GCTGC**CAGACAAAGAGAACGCAGCACGA**CGTAG**CG
ACGCAGACGGCGCAGCGAG**CATAG**CACGCACGCAGCCACGCACAGAC**CGTCG**CCAGCC
GCAGCAGCACGACA**CATCG**CGACGGCACGGAGCGGACGCACGACGAGCACAAAACAAAA
AAAAAAAA

2′-Deoxyinosine (I) containing DNA coding strand

5′- AAG CTA ATA CGA CTC ACT ATA GGA GCA GCA CGA CGA ACG GAC GAG GTI CCA
CGA CAG AGA GCG GAC GCA ATI CCG ACC GAC GAA CAC GCA GCT ICC AGA CAA AGA
GAA CGC AGC ACG ACI TGC CGA CGC AGA CGG CGC AGC GAG CIT ACC ACG CAC GCA
GCC ACG CAC AGA CGI TCC ACA GCC GCA GCA GCA CGA ACG ACG ACA AAA CAA AAA
AAA AAA A

The IVT experiments were conducted with duplex DNA comprised of the coding strands shown annealed with their complementary strands. For the I-containing DNA coding strand, the complementary template strand had a dC nucleotide placed opposite.

**Figure S2.** Tombo analysis to compare the current levels between U and Ψ or U and m$^1$Ψ.

The Tombo plots are grouped to also show Ψ and m$^1$Ψ in the same sequence context together. The arrow in each plot is the site with U, Ψ, or m$^1$Ψ

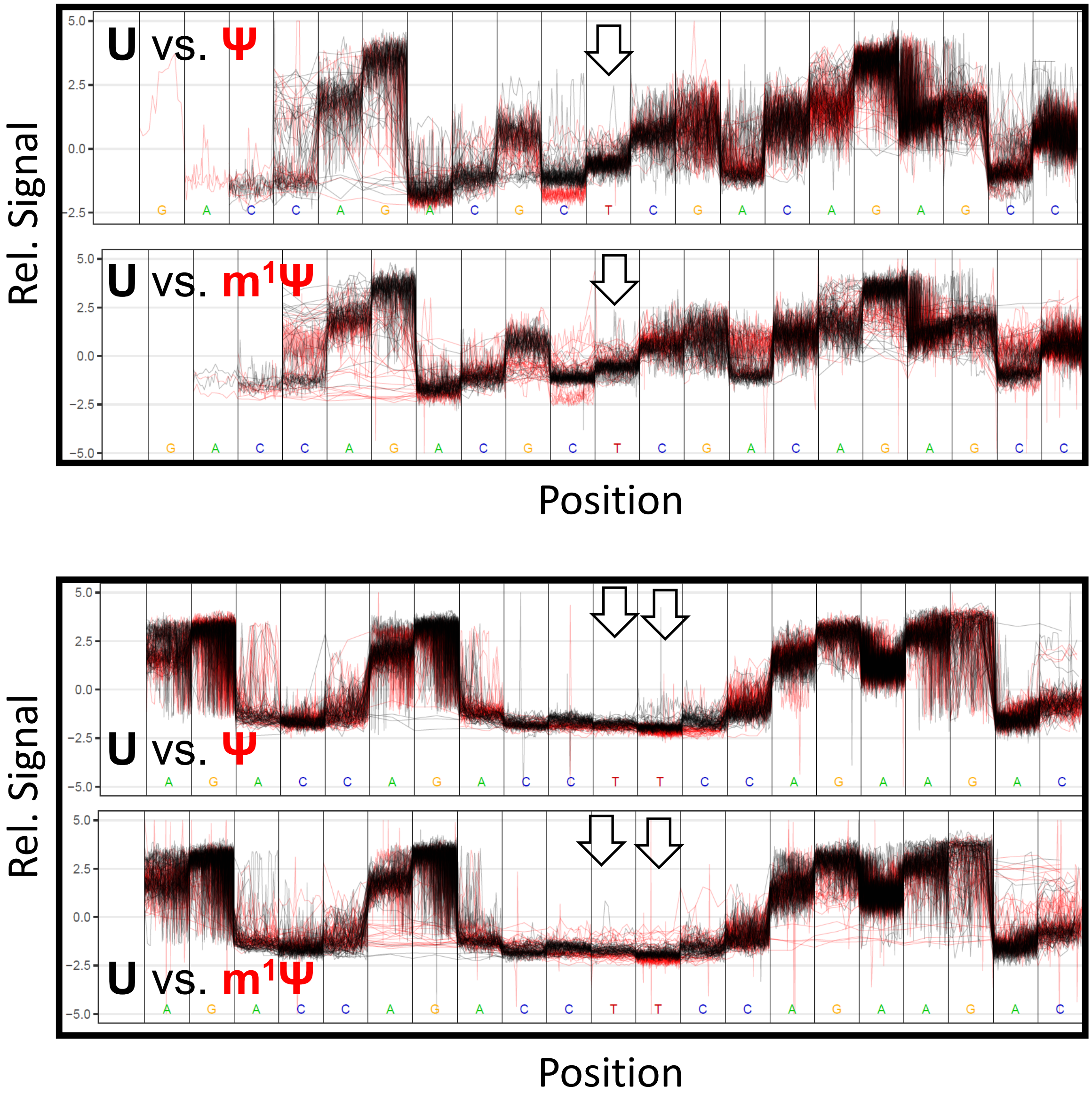

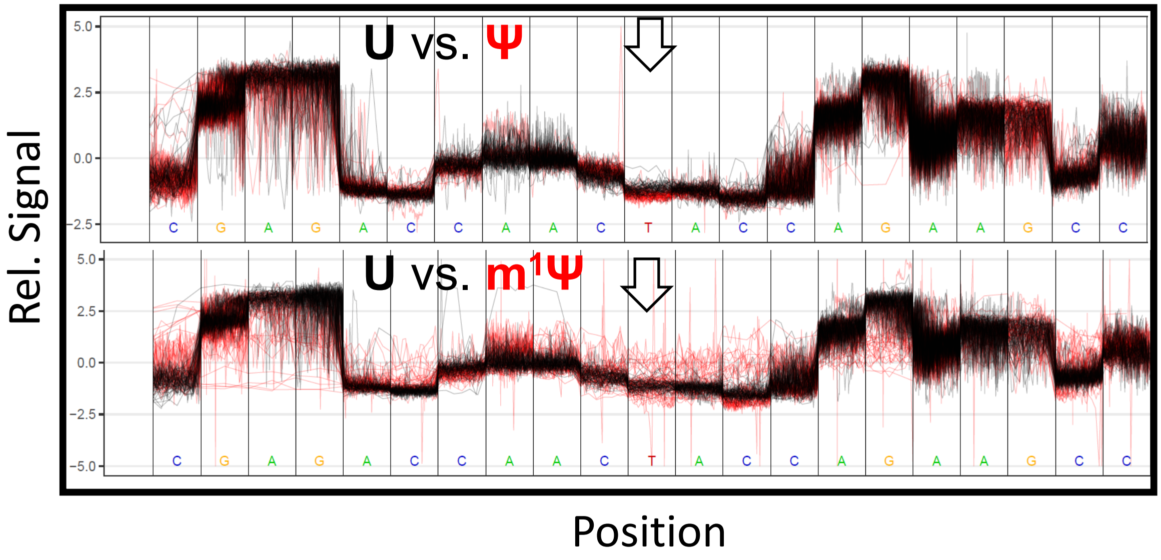
U vs. Ψ
U vs. m1Ψ
Rel. Signal
Position
C G A G A C C A A C T A C C A G A A G C C

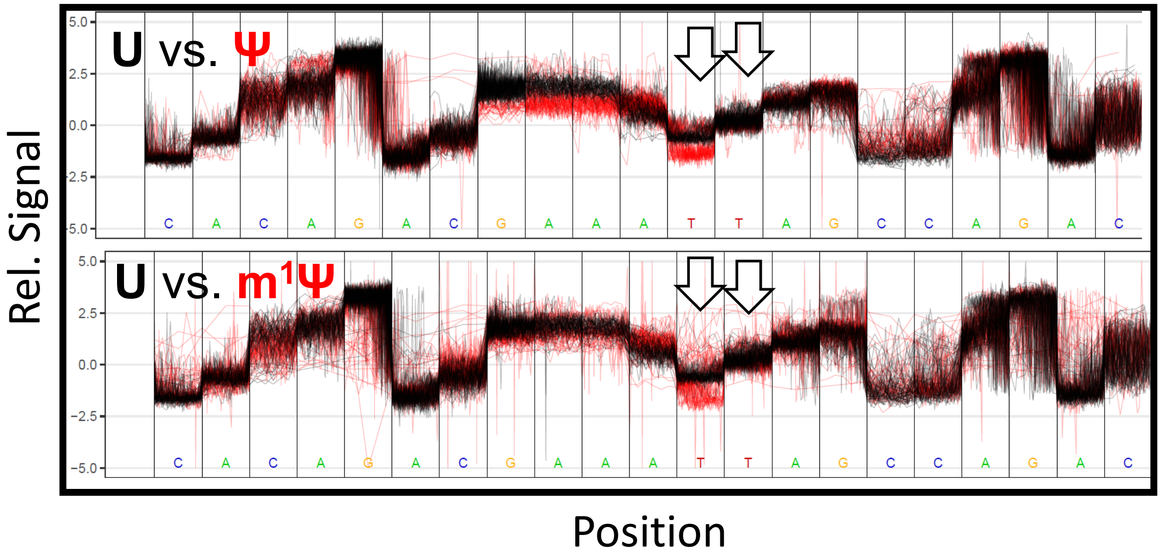
U vs. Ψ
U vs. m1Ψ
Rel. Signal
Position
C A C A G A C G A A A T T A G C C A G A C

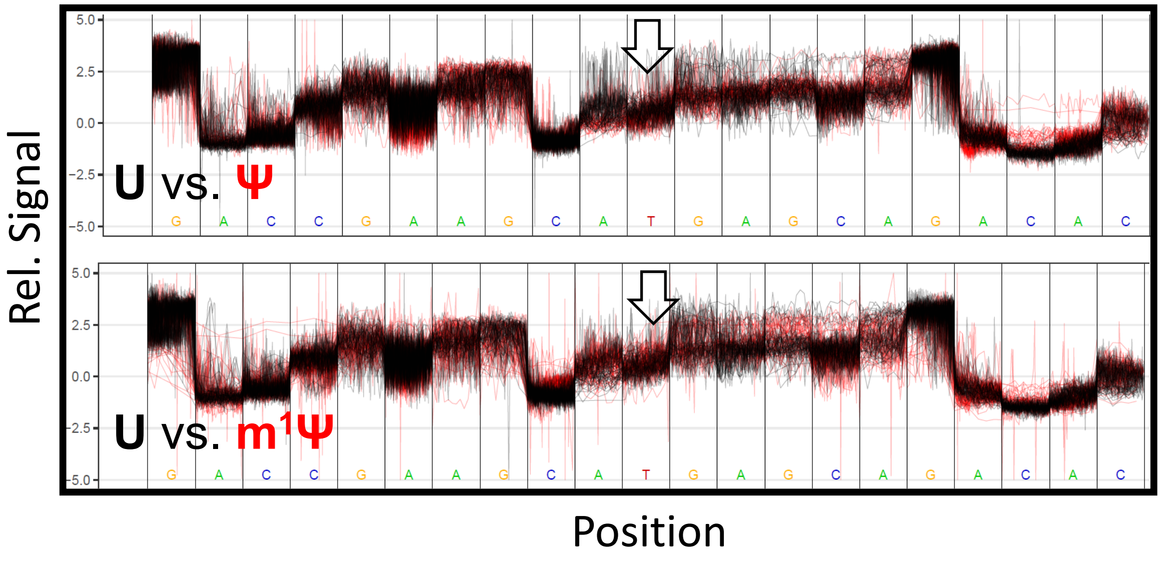
Rel. Signal
5.0
2.5
0.0
-2.5
-5.0
U vs. Ψ
G A C C G A A G C A T G A G C A G A C A C
U vs. m1Ψ
G A C C G A A G C A T G A G C A G A C A C
Position

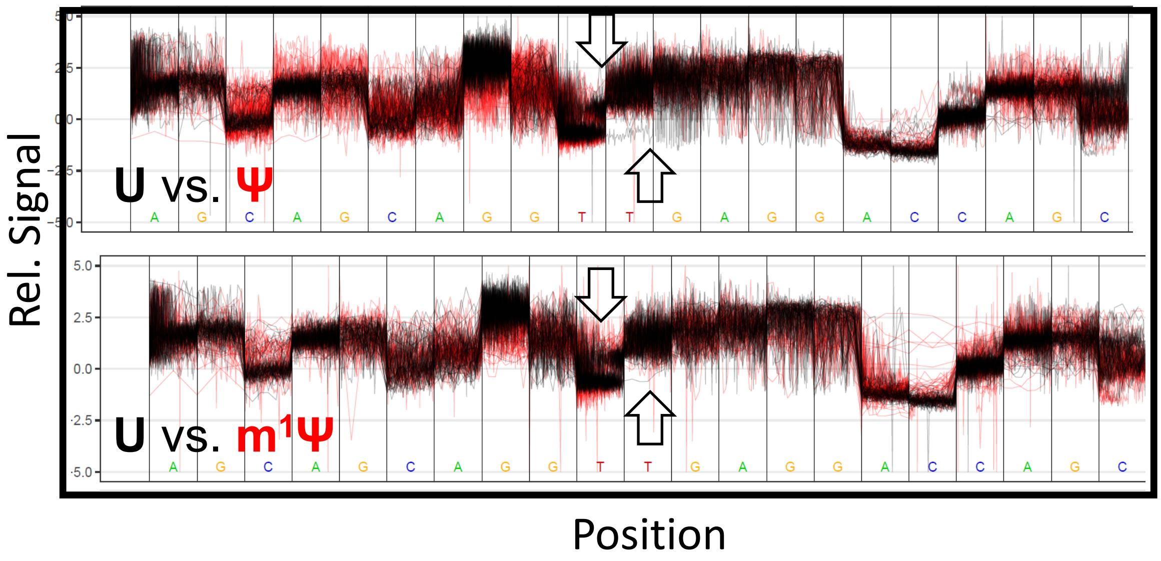
Rel. Signal
5.0
2.5
0.0
-2.5
-5.0
U vs. Ψ
A G C A G C A G G T T G A G G A C C A G C
U vs. m1Ψ
A G C A G C A G G T T G A G G A C C A G C
Position

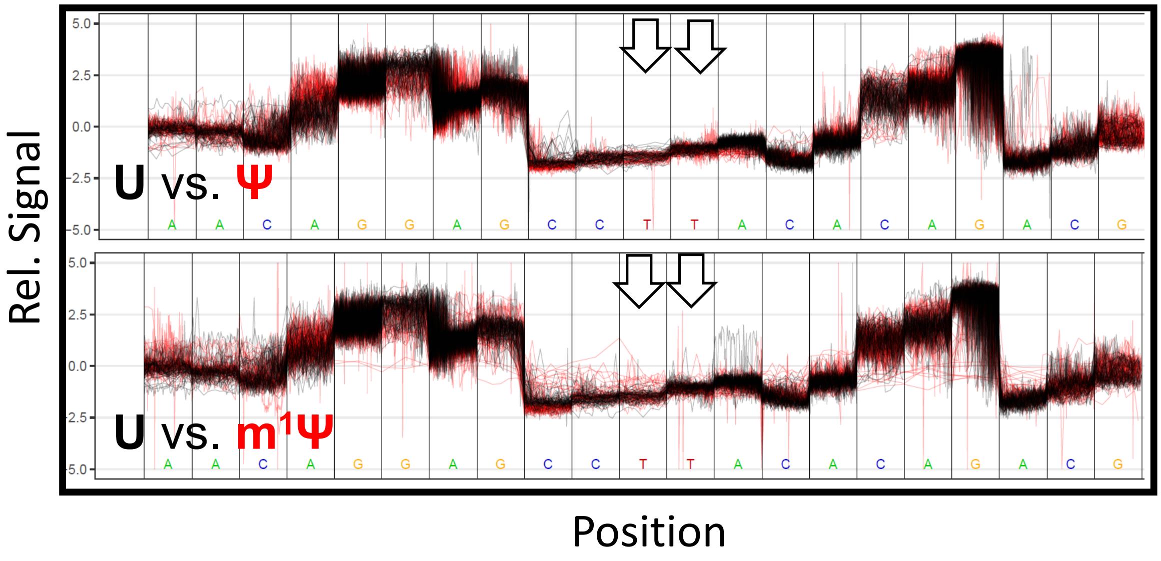
Rel. Signal
U vs. Ψ
U vs. m1Ψ
5.0
2.5
0.0
-2.5
-5.0
A A C A G G A G C C T T A C A C A G A C G
Position

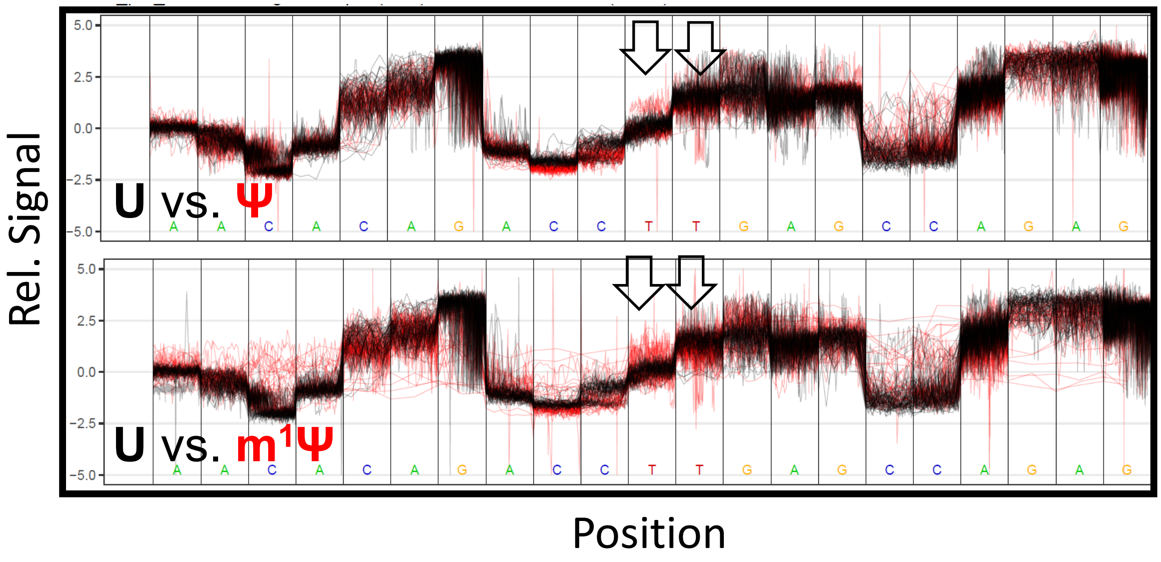
Rel. Signal
U vs. Ψ
U vs. m1Ψ
5.0
2.5
0.0
-2.5
-5.0
A A C A C A G A C C T T G A G C C A G A G
Position

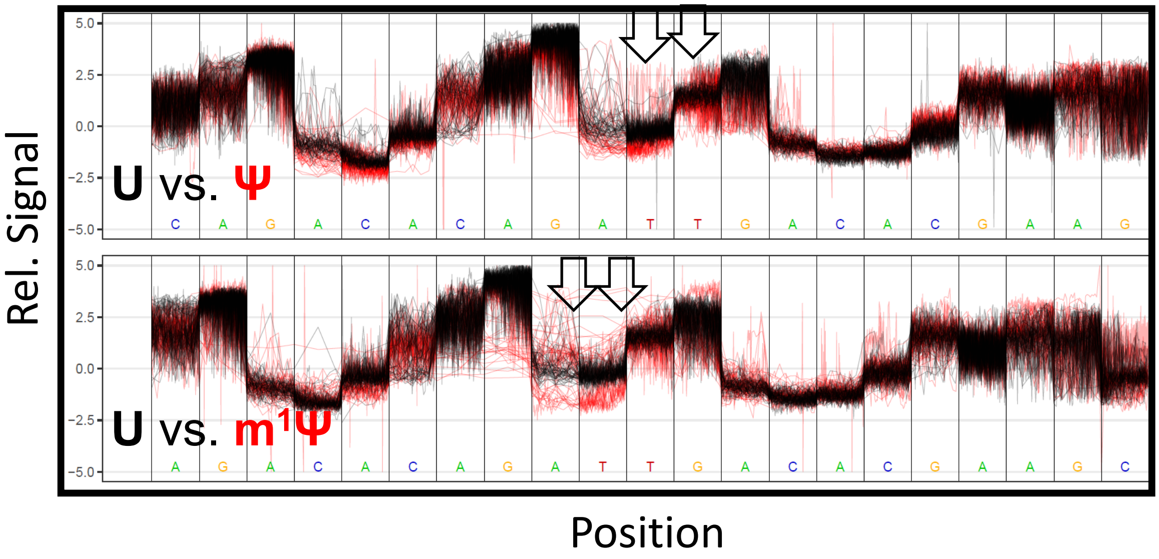
Rel. Signal
5.0
2.5
0.0
-2.5
-5.0
U vs. Ψ
C A G A C A C A G A T T G A C A C G A A G
U vs. m1Ψ
A G A C A C A G A T T G A C A C G A A G C
Position

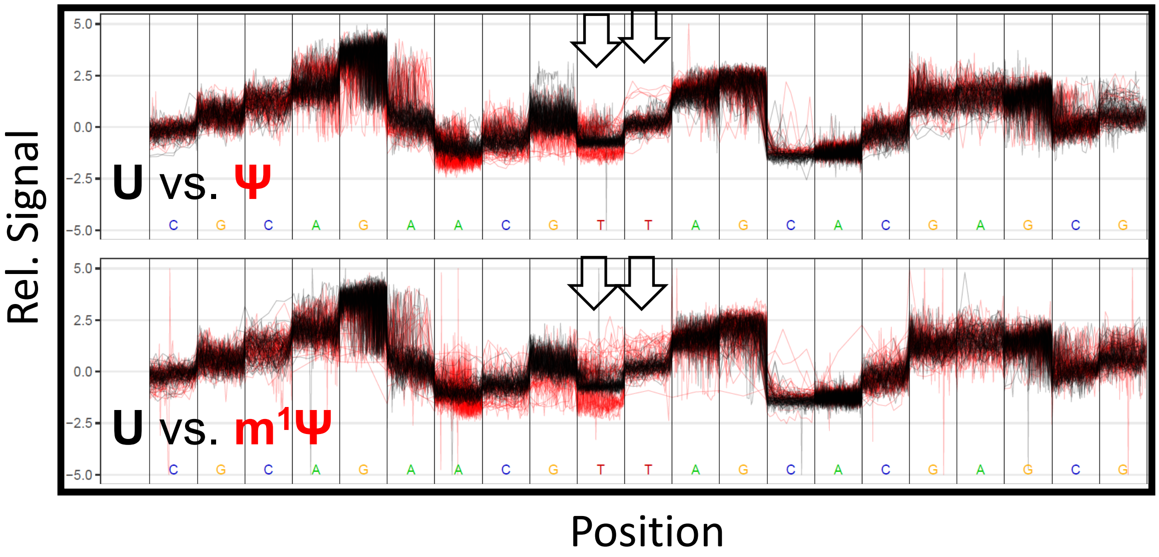
Rel. Signal
5.0
2.5
0.0
-2.5
-5.0
U vs. Ψ
C G C A G A A C G T T A G C A C G A G C G
U vs. m1Ψ
C G C A G A A C G T T A G C A C G A G C G
Position

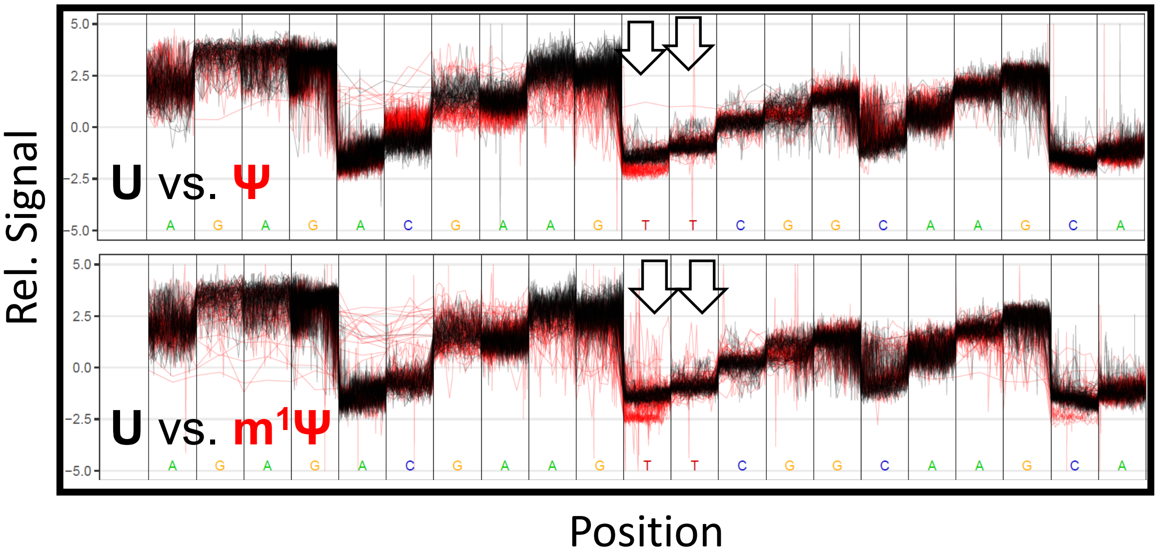
Rel. Signal
5.0
2.5
0.0
-2.5
-5.0
U vs. Ψ
U vs. m1Ψ
A G A G A C G A A G T T C G G C A A G C A
Position

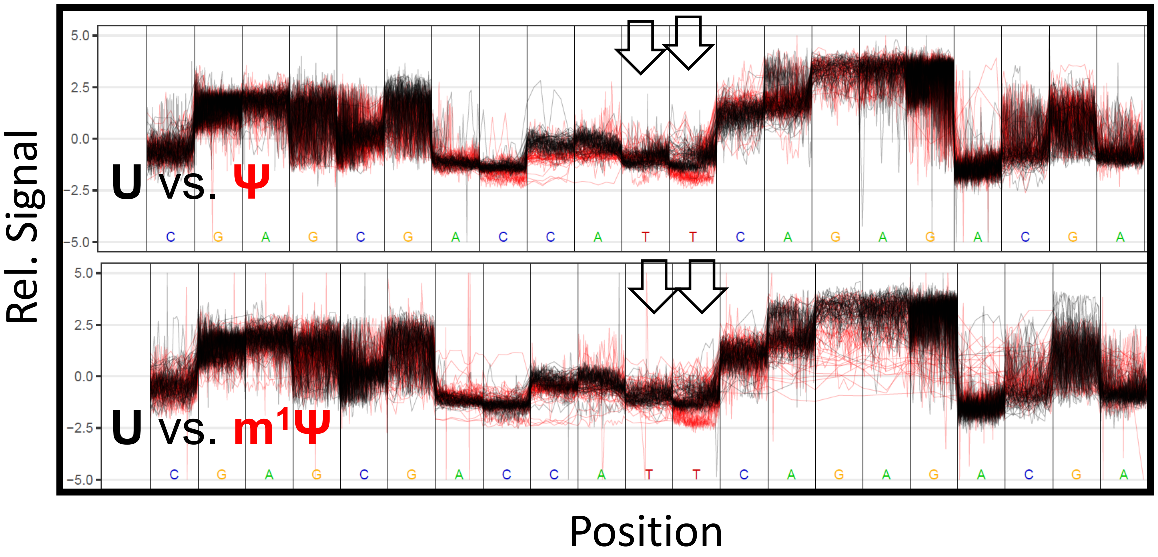
Rel. Signal
5.0
2.5
0.0
-2.5
-5.0
U vs. Ψ
U vs. m1Ψ
C G A G C G A C C A T T C A G A G A C G A
Position

**Figure S3.** Percent reads recovered after Nanopolish resquiggling.

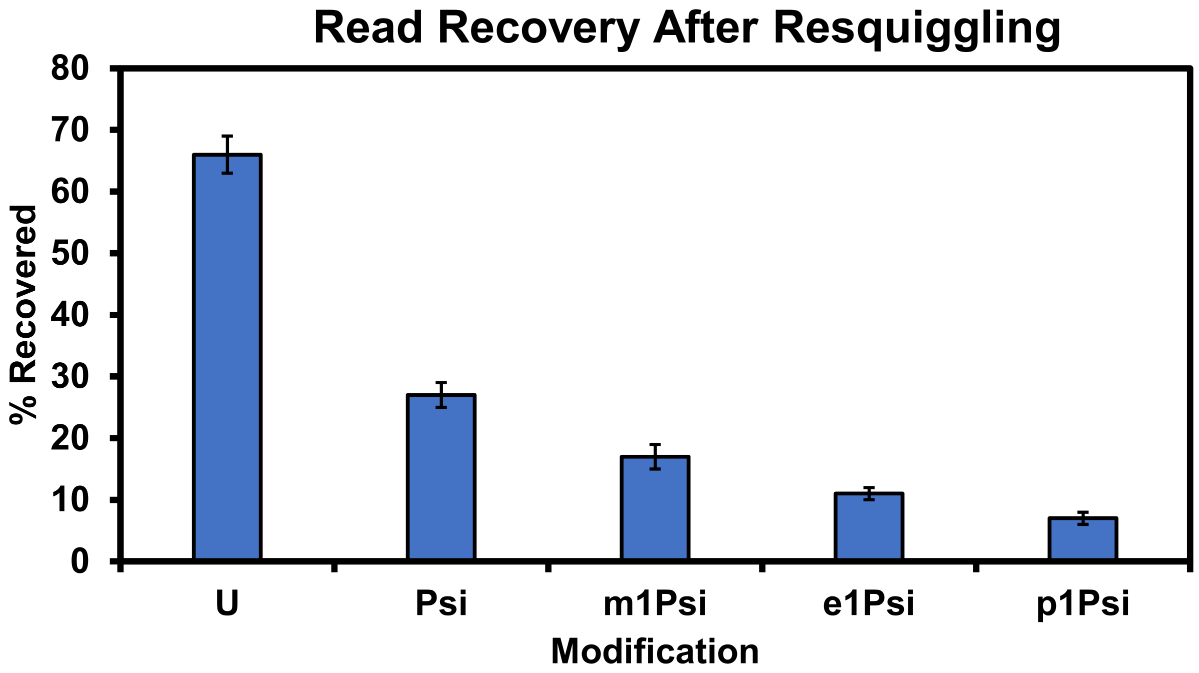

**Figure S4.** Spearman's rank-order correlation test on base call error vs. current levels and dwell times.

U vs. Ψ correlation analysis

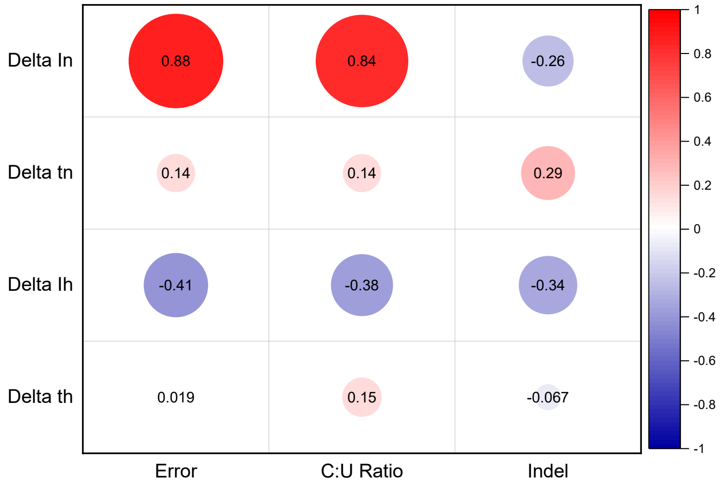

U vs. m[1]Ψ correlation analysis

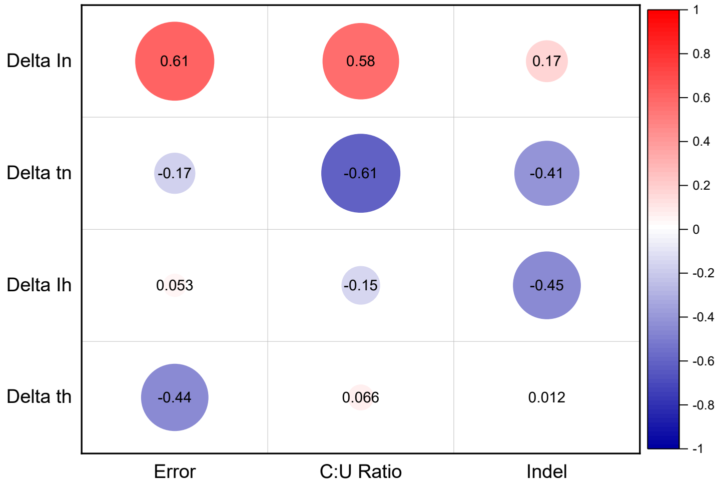

**Figure S5.** Additional nanopore current and dwell data for the N1-alkylpseudouridine derivatives.

| Sequence Context | Nucleotide (X) | Nanopore Current $I_N$ (pA) | Nanopore Dwell Time $t_N$ (msec) | Helicase Current $I_H$ (pA) | Helicase Dwell Time $t_H$ (msec) |
|---|---|---|---|---|---|
| 5`-GGXGA | U | 120.1 +- 5.7 | 0.012 +- 0.01 | 120.9 +- 8.0 | 0.01 +- 0.005 |
| | Pseudo-U | 110.3 +- 4.6 | 0.012 +- 0.01 | 120.6 +- 7.6 | 0.026 +- 0.01 |
| | N1-Methylpseudo-U | 110.7 +- 5.7 | 0.011 +- 0.01 | 120.1 +- 7.7 | 0.03 +- 0.01 |
| | N1-Ethylpseudo-U | 110.5 +- 5.7 | 0.011 +- 0.01 | 121.2 +- 7.9 | 0.028 +- 0.01 |
| | N1-Propylpseudo-U | 109.7 +- 2.4 | 0.011 +- 0.01 | 121.7 +- 5.6 | 0.033 +- 0.01 |
| 5`-AGXGA | U | 78.1 1.8 | 0.012 +- 0.006 | 85.3 +- 3 | 0.01 +- 0.005 |
| | Pseudo-U | 74.1 +- 2.1 | 0.011 +- 0.008 | 86.4 +- 4.1 | 0.0232 +- 0.01 |
| | N1-Methylpseudo-U | 73.2 +- 1.8 | 0.011 +- 0.006 | 86.0 +- 3.1 | 0.03 +- 0.01 |
| | N1-Ethylpseudo-U | 71.2 +- 3.2 | 0.013 +- 0.005 | 86.1 +- 3.1 | 0.045 +- 0.01 |
| | N1-Propylpseudo-U | 70.0 +- 3.7 | 0.014 +- 0.007 | 87.0 +- 2.3 | 0.046 +- 0.01 |
| 5`-GCXGC | U | 83.8 +- 2.1 | 0.012 +- 0.009 | 138.4 +- 4.7 | 0.02 +- 0.008 |
| | Pseudo-U | 85.8 +- 1.5 | 0.018 +- 0.004 | 138.3 +- 3.9 | 0.04 +- 0.01 |
| | N1-Methylpseudo-U | 80.8 +- 2.1 | 0.012 +- 0.009 | 138.7 +- 4.7 | 0.03 +- 0.01 |
| | N1-Ethylpseudo-U | 78.6 +- 3.9 | 0.011 +- 0.009 | 139.2 +- 4.8 | 0.04 +- 0.01 |
| | N1-Propylpseudo-U | 78.1 +- 2.5 | 0.013 +- 0.009 | 138.4 +- 4.1 | 0.044 +- 0.01 |
| 5`-CGXAG | U | 102.1 +- 4.2 | 0.017 +- 0.005 | 131.6 +- 3.6 | 0.023 +- 0.005 |
| | Pseudo-U | 97.9 +- 4.1 | 0.01 +- 0.005 | 132.0 +- 3.5 | 0.029 +- 0.008 |
| | N1-Methylpseudo-U | 96.1 +- 3.7 | 0.015 +- 0.009 | 132.6 +- 3.7 | 0.031 +- 0.005 |
| | N1-Ethylpseudo-U | 96.2 +- 5.3 | 0.014 +- 0.007 | 133.0 +- 4.1 | 0.030 +- 0.005 |
| | N1-Propylpseudo-U | 95.9 +- 4.1 | 0.015 +- 0.005 | 132.7 +- 4.6 | 0.03 +- 0.01 |
| 5`-CAXAG | U | 90.1 +- 3.1 | 0.01 +- 0.005 | 94.5 +- 4.9 | 0.01 +- 0.008 |
| | Pseudo-U | 86.0 +- 3.1 | 0.01 +- 0.005 | 94.4 +- 5.3 | 0.016 +- 0.007 |
| | N1-Methylpseudo-U | 85.9 +- 3.0 | 0.01 +- 0.005 | 93.9 +- 6.3 | 0.015 +- 0.008 |
| | N1-Ethylpseudo-U | 83.0 +- 3.1 | 0.01 +- 0.005 | 94.2 +- 4.7 | 0.017 +- 0.007 |
| | N1-Propylpseudo-U | 76.2 +- 3.1 | 0.015 +- 0.006 | 93.9 +- 4.6 | 0.018 +- 0.005 |
| 5`-CGXCG | U | 93 +- 2.7 | 0.016 +- 0.006 | 98.2 +- 3.6 | 0.02 +- 0.01 |
| | Pseudo-U | 95.1 +- 2.9 | 0.018 +- 0.007 | 97.9 +- 2.6 | 0.05 +- 0.009 |
| | N1-Methylpseudo-U | 92.4 +- 2.7 | 0.016 +- 0.005 | 98.5 +- 3.5 | 0.04 +- 0.01 |
| | N1-Ethylpseudo-U | 90.5 +- 2.8 | 0.011 +- 0.005 | 98.5 +- 3.2 | 0.04 +- 0.01 |
| | N1-Propylpseudo-U | 81.9 +- 2.6 | 0.013 +- 0.006 | 98.3 +- 3.7 | 0.045 +- 0.01 |
| 5`-CAXCG | U | 81.2 +- 2.9 | 0.013 +- 0.007 | 82.1 +- 3.8 | 0.017 +- 0.008 |
| | Pseudo-U | 79.9 +- 2.1 | 0.015 +- 0.006 | 84.5 +- 2.9 | 0.025 +- 0.003 |
| | N1-Methylpseudo-U | 79.4 +- 2.2 | 0.013 +- 0.007 | 82.0 +- 2.7 | 0.027 +- 0.005 |
| | N1-Ethylpseudo-U | 77.6 +- 2.3 | 0.02 +- 0.01 | 81.7 +- 2.1 | 0.053 +- 0.007 |
| | N1-Propylpseudo-U | 76.2 +- 3.8 | 0.013 +- 0.007 | 81.9 +- 2.6 | 0.033 +- 0.011 |

The values were obtained from Nanopolish analysis.

**Figure S6.** Full-length extension evaluated by agarose gel electrophoresis.

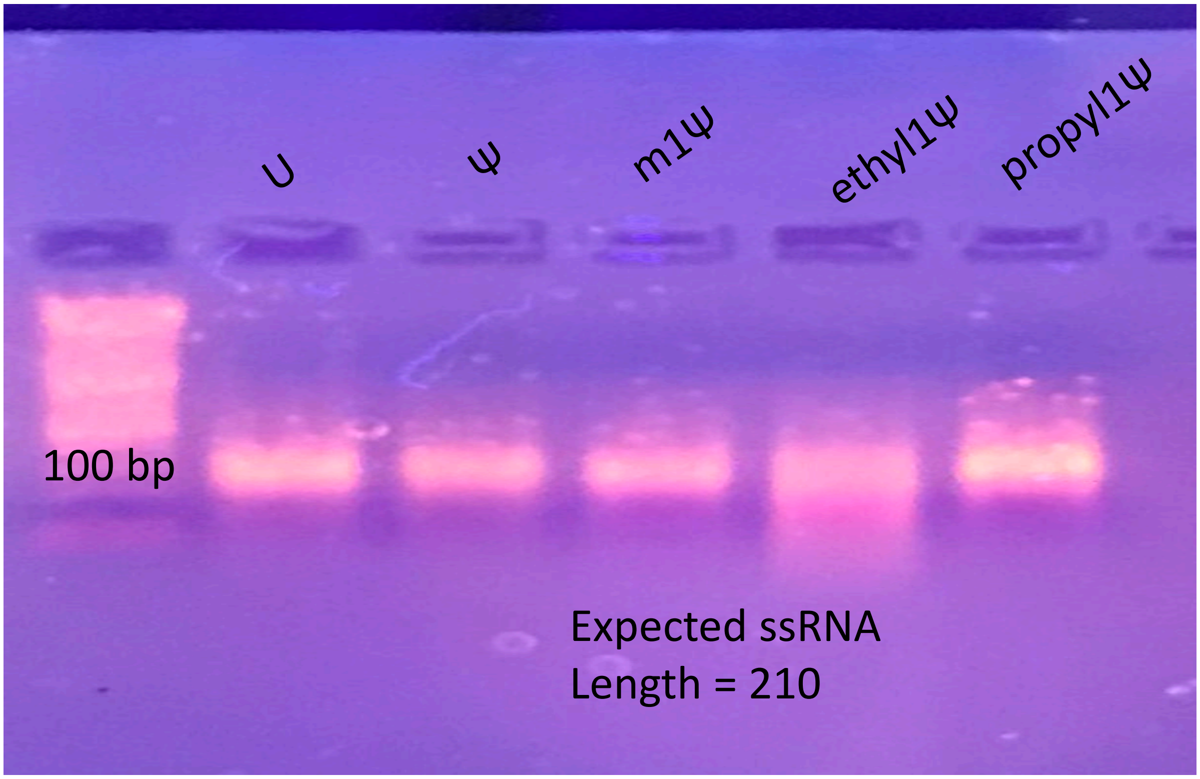

**Figure S7.** Calibration curves to quantify Ψ or $m^1Ψ$ developed with Nanopore-Psu. Provided are examples of calibration curves generated.

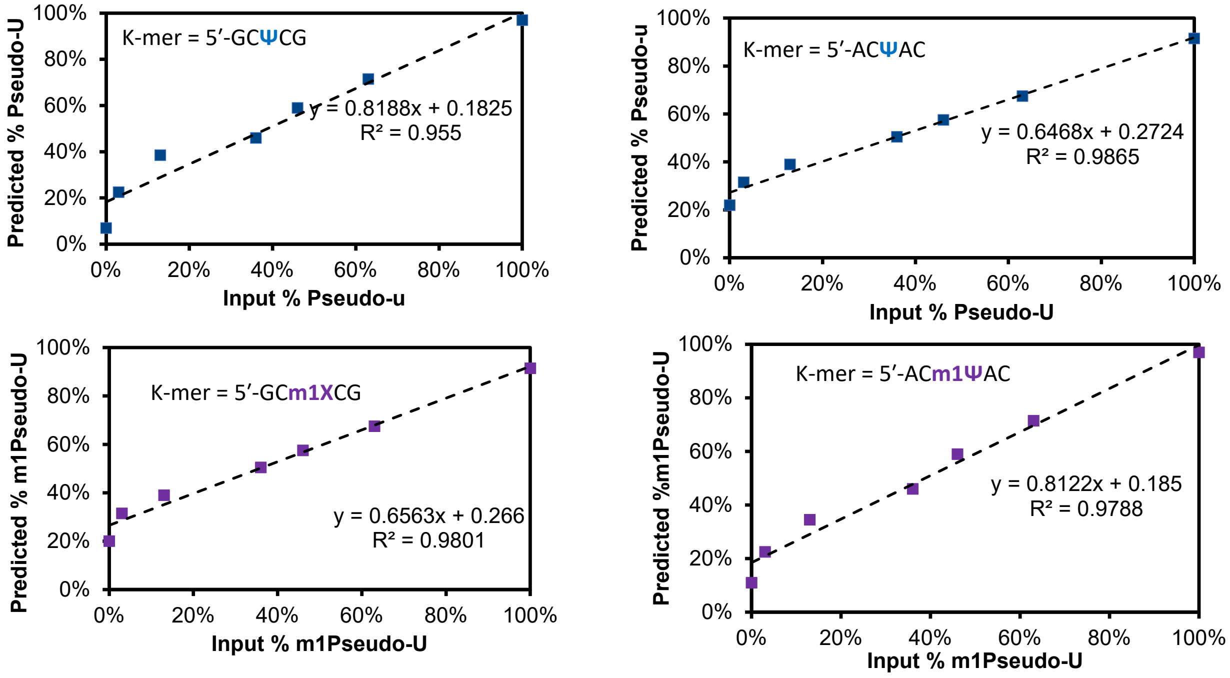

**Figure S8.** Controls with a dI:dC base pair on the 5′ side of the T7 RNA polymerase competition site.

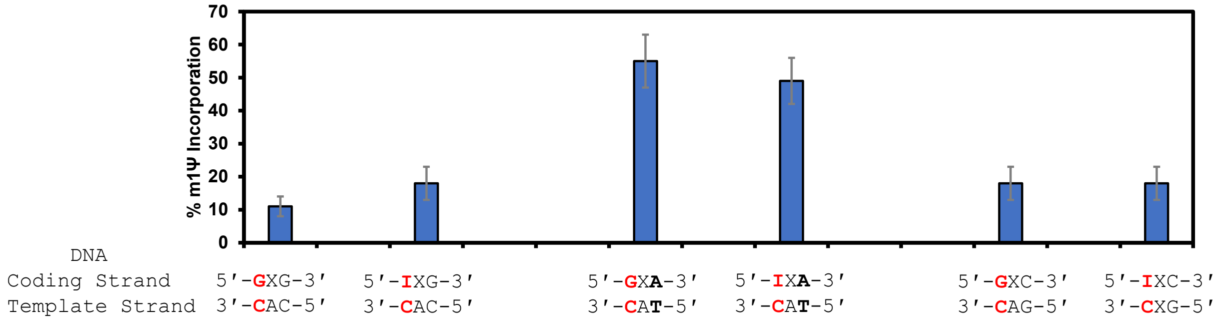

**Figure S9.** T7 RNA polymerase studies that competed UTP vs. e[1]ΨTP or p[1]ΨTP for insertion and elongation.

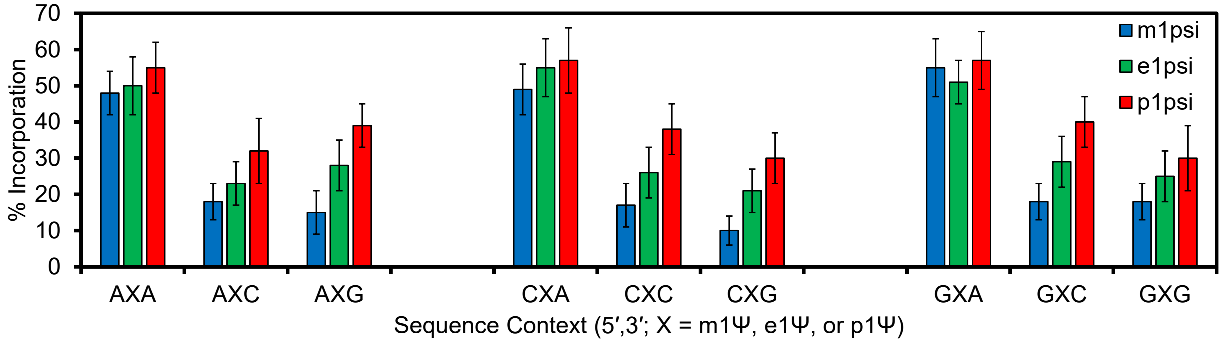
